# Comparative phylogenomics reveals multilevel genomic differentiation between annual and perennial *Hordeum* species

**DOI:** 10.1101/2025.08.11.669716

**Authors:** Timo Hellwig, Nina Döring, Einar Baldvin Haraldsson, Maria von Korff

## Abstract

Annual plants complete their life cycle with a single reproductive episode, whereas perennials must balance reproduction with the maintenance of tissues that sustain future growth. Despite the fundamental importance of this distinction, the genomic differences associated with these contrasting life histories remain poorly understood. Here, we utilize independent occurrences of annuality in the grass genus *Hordeum* to identify recurrent differences between annual and perennial species. Combining transcriptomes from 82 accessions with recently published reference genomes, we compared protein-coding sequence evolution, gene copy number, gene expression and co-expression while accounting for evolutionary non-independence. We found widespread genomic differentiation, with several consistent patterns. Genes involved in cell division and genome maintenance showed evidence of stronger evolutionary constraint in perennials, while gene families associated with developmental regulation and senescence were more frequently retained in higher copy numbers in perennials. Differences in gene expression and coordination were particularly pronounced during reproduction, when perennials showed higher expression of genes involved in cell division, cytoskeletal organisation, cell-wall formation and resource transport, together with extensive reorganisation of gene co-expression relationships. Across these complementary analyses, processes involved in cellular proliferation, developmental regulation and tissue maintenance repeatedly distinguished the two life histories. Our findings suggest that annual-perennial differentiation extends beyond individual flowering regulators to broader genomic differences associated with a more complex coordination of processes balancing growth and senescence in perennials. By identifying these recurrent patterns across evolutionary lineages, this study provides a comparative framework for understanding the genomic differentiation underlying contrasting plant life histories.

## Introduction

Plants exhibit remarkable diversity in life-history strategies to maximise survival and reproduction in specific environments. A fundamental distinction is between annual and perennial plants. Annuals complete their life cycle within one growing season and are semelparous (monocarpic), dying after reproduction. Perennials live for multiple years, ranging from biennials with one vegetative and one reproductive year to woody perennials surviving for centuries and reproducing repeatedly (Friedman 2020). These alternative life-history strategies are accompanied by broader differences in growth and resource allocation. Annuals generally develop and reproduce more rapidly and allocate more strongly to reproduction, whereas perennials must balance current reproduction against the maintenance of tissues and meristems required for future growth and reproduction (Garnier 1992; Lundgren and Des Marais 2020; Hjertaas et al., 2023; Anokye et al., 2026). Despite these substantial developmental and physiological differences, transitions between annual and perennial life histories have occurred repeatedly across angiosperms, indicating that life-history strategy is evolutionarily labile (Friedman 2020; Hjertaas et al., 2023).

Understanding the basis and consequences of contrasting life-history strategies is therefore a central question in plant evolutionary biology. This question has recently gained particular relevance in agriculture, where perennial crops can provide permanent soil cover that reduces erosion (Vallebona et al., 2016; Li et al., 2022) and deeper root systems that minimise nutrient leaching (Crews and Rumsey 2017; Lundgren and Des Marais 2020), and are increasingly recognised as a component of more sustainable agricultural systems (Crews et al., 2018; Zhao and Wang 2024). Several approaches have therefore aimed to develop perennial grain crops (Chapman et al., 2022; Zhao and Wang 2024), either through domestication of perennial species, as exemplified by intermediate wheatgrass (DeHaan et al., 2020), or through hybridisation between annual crops and perennial relatives, as in perennial rice (Zhang et al., 2023). Yet combining perennial persistence with the reproductive output and agronomic performance of annual crops remains difficult (Chapman et al., 2022), highlighting how incompletely the biological basis of perennial growth is still understood. This gap extends beyond crop development: despite the repeated evolution of annual and perennial life histories across angiosperms, their underlying genetic basis has been investigated in only a small number of model systems.

The best-characterised examples indicate that different developmental mechanisms can contribute to annual-perennial differentiation. In Brassicaceae, flowering-time and meristem regulators play prominent roles in controlling seasonal flowering and the maintenance of perennial growth, and experimental manipulation of a small number of such regulators can shift plants towards annual or perennial-like growth forms (Bastow et al., 2004; Sung and Amasino 2004; Melzer et al., 2008; Wang et al., 2009; Zhai et al., 2024). In hybrid as-pen, seasonal growth and dormancy are instead strongly regulated by environmental cues such as photoperiod (Böhlenius et al., 2006). Genetic studies in grasses remain limited and have largely focused on traits associated with vegetative persistence, particularly rhizome formation in perennial cereals and their wild relatives (Hu et al., 2003; Hu et al., 2011; Gruner and Miedaner 2021; Zhang et al., 2023). Thus, existing studies have identified important components of perennial growth in individual systems, but provide a limited basis for determining which genomic differences recur across independently evolved annual and perennial lineages.

The genus *Hordeum* provides a suitable system for addressing this question. *Hordeum* includes both annual and perennial species, with at least three independent transitions between life-history strategies (Brassac and Blattner 2015). Most perennial *Hordeum* species are iteroparous, maintaining living above-ground vegetative shoots through winter with reduced growth and reinitiating reproductive growth under long days. A notable exception is the Mediterranean species H. bulbosum, which dies above ground after seed set and survives the unfavourable Mediterranean summer as dormant underground bulbs that resprout after autumn rains (Koller and Highkin 1960; Fuerst et al., 2023). Although such pseudo-annual species are perennial in the genetic sense because the same genotype persists across years, their persistence resembles that of annuals ecologically: the unfavourable season is bridged by a dormant propagule rather than by continuously maintained above-ground shoots and meristems. Herein, we therefore operationally define perennial *Hordeum* species according to ecological persistence strategy rather than persistence of the genotype itself and exclude *H. bulbosum* from the main annual-perennial comparisons.

Repeated independent occurrences of traits such as annual-perennial life-history strategies provide the evolutionary replication needed to distinguish trait-associated patterns from clade-specific divergence. In *Hordeum*, the occurrence of annuality in phylogenetically separated lineages provides exactly such a comparative setting. Phylogenetically informed statistical models can further strengthen these comparisons by explicitly accounting for the non-independence among accessions and species caused by shared ancestry. Their application, however, depends on a well-resolved evolutionary framework, because incomplete lineage sorting and hybridisation can obscure species relationships and complicate comparative inference (Soltis and Soltis 2003). This is particularly relevant in *Hordeum*, where relationships among some recently diverged lineages have remained uncertain and interspecific hybridisation has contributed to the evolutionary history of the genus. The recent publication of genomes for all 21 diploid and three tetraploid *Hordeum* species (Feng et al., 2025) enables comparative genetic studies of the evolution and genetic architecture of various traits, including life-history strategies in the *Hordeum* clade.

Here, we combined newly generated tissue and stage specific transcriptomes from multiple accessions across *Hordeum* with recently published reference genomes to identify genomic features repeatedly associated with annual and perennial life histories. We first reconstructed accession- and species-level relationships across the genus and investigated interspecies introgression, establishing the evolutionary framework required for phylogenetically informed comparisons. We then examined annual-perennial differentiation at four complementary genomic levels: selective constraint on protein-coding genes, gene copy-number variation, mean gene expression and gene co-expression. By integrating these analyses across phylogenetically separated occurrences of annuality, we asked which genomic features consistently distinguish annual and perennial *Hordeum* species, and whether different levels of genomic variation reveal recurrent biological processes associated with life-history differentiation.

## Results & Discussion

### *Hordeum* phylogeny and interspecies hybridization

To identify genomic features repeatedly associated with annual and perennial life histories, we first established a phylogenetic framework that resolves both species and among-accession relationships across *Hordeum*. This was necessary because our downstream analyses used different data types and levels of biological replication: accession-level relationships were needed for phylogenetically informed expression and selection analyses, whereas species-level relationships were required for analysing gene copy-number variation. We therefore inferred three complementary phylogenies and asked whether they converged on the same overall evolutionary structure. Because evolutionary relationships may also include reticulate histories that cannot be adequately represented by strictly bifurcating trees, we complemented these reconstructions with an analysis of interspecies introgression to provide a more comprehensive reconstruction of *Hordeum* evolutionary history.

First, we generated a maximum-likelihood (ML) accession-based tree using the combined RNA-seq and reference-genome dataset. Each tip represented an individual accession, allowing us to capture intraspecific variation and evaluate the robustness of species boundaries. This tree was based on 257 single-copy ortho logs (scHOGs) shared across all *Hordeum* and outgroup accessions, with a total alignment length of 440,610 bp and 52,026 parsimony-informative sites (Tab. S1). Second, we constructed a Bayesian species tree under the multispecies coalescent while simultaneously estimating divergence times. Because of its substantially higher computational demands, this analysis was restricted to 90 scHOGs, comprising 225,942 bp and 22,571 parsimony-informative sites (Tab. S1). Third, we constructed an ML tree using only the reference genomes from Feng et al. (2025). This genome-accession tree was primarily generated for visualisation of gene copy-number variation and contained one accession per species, together with several more distant out - groups. It was based on 380 scHOGs, with a total alignment length of 558,769 bp and 115,640 parsimony-in-formative sites (Tab. S1).

The topologies of the three trees were almost identical (Fig. 1, 3, S1, S2). The *Hordeum* clade split into two major monophyletic groups approximately 10.3 million years ago (95% HPD: 8.2–15.8 million years; Fig. S3). One comprised sections *Hordeum* and Trichostachys, containing species distributed across Europe, the Mediterranean and Central Asia. All species in these sections are annual except the pseudo-annual *H. bulbo-sum* (Fig. 1, S2). The second major group comprised sections Marina and Stenostachys. Section Marina contained two annual species from Europe and the Mediterranean, whereas Stenostachys comprised the Asian series Sibirica and the American series Critesion. Most Stenostachys species were perennial, with annuality reappearing among the three most recently diverged species, H. pusillum, H. euclaston and H. intercedens. Annual species therefore occur in two phylogenetically separate parts of the tree, allowing annual-perennial differences to be distinguished from clade-specific effects of shared ancestry.

**Figure 1:**
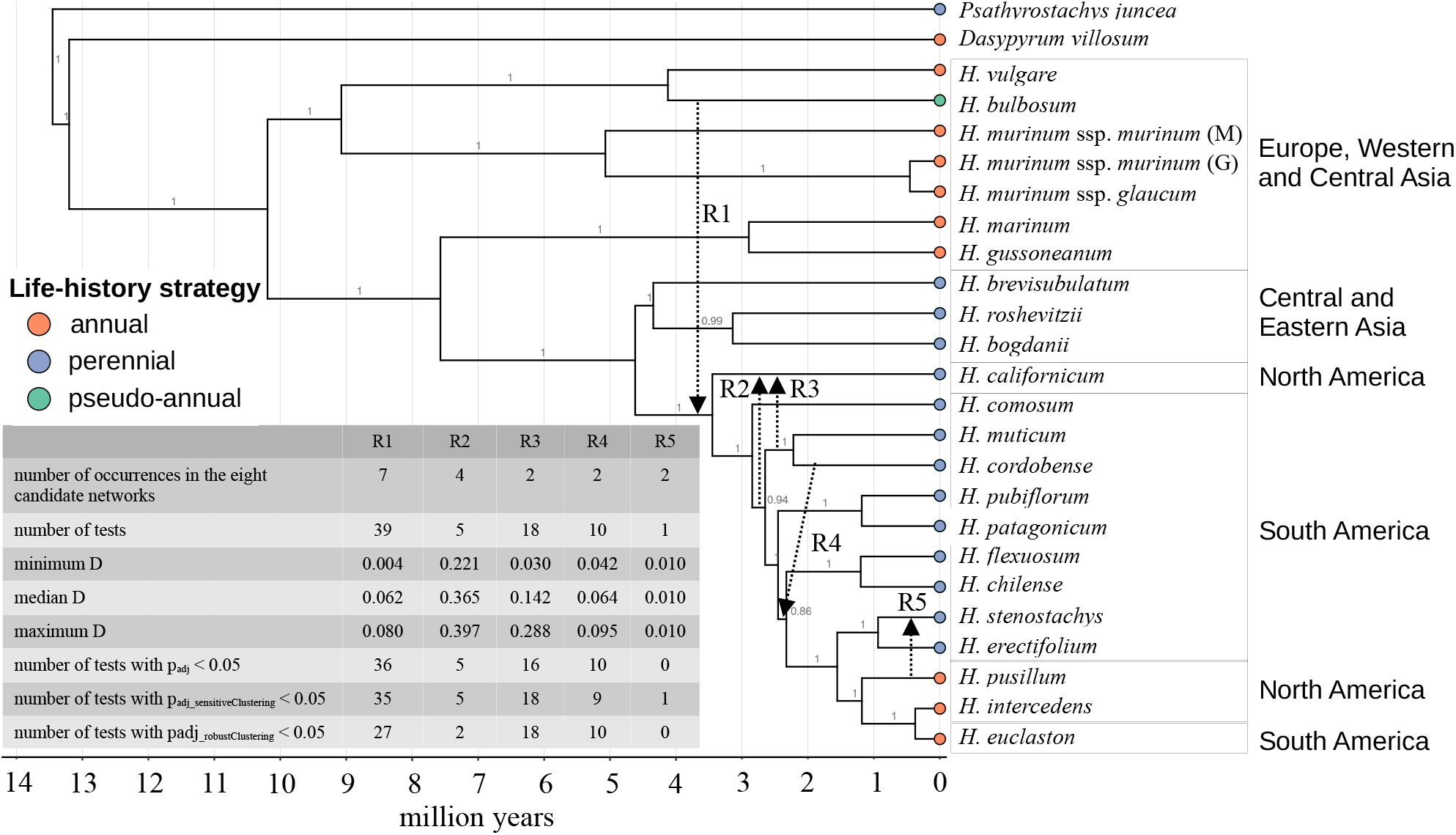
Bayesian species tree with inter-species introgression events. Numbers at the nodes are posterior probabilities. Species names are coloured according to their geographic distribution. Tip colours represent life-cycle strategy. Lines R1-5 represent candidate reticulations. The table summarize the results of the ABBA-BABA tests (Tab. S2).

**Figure 2:**
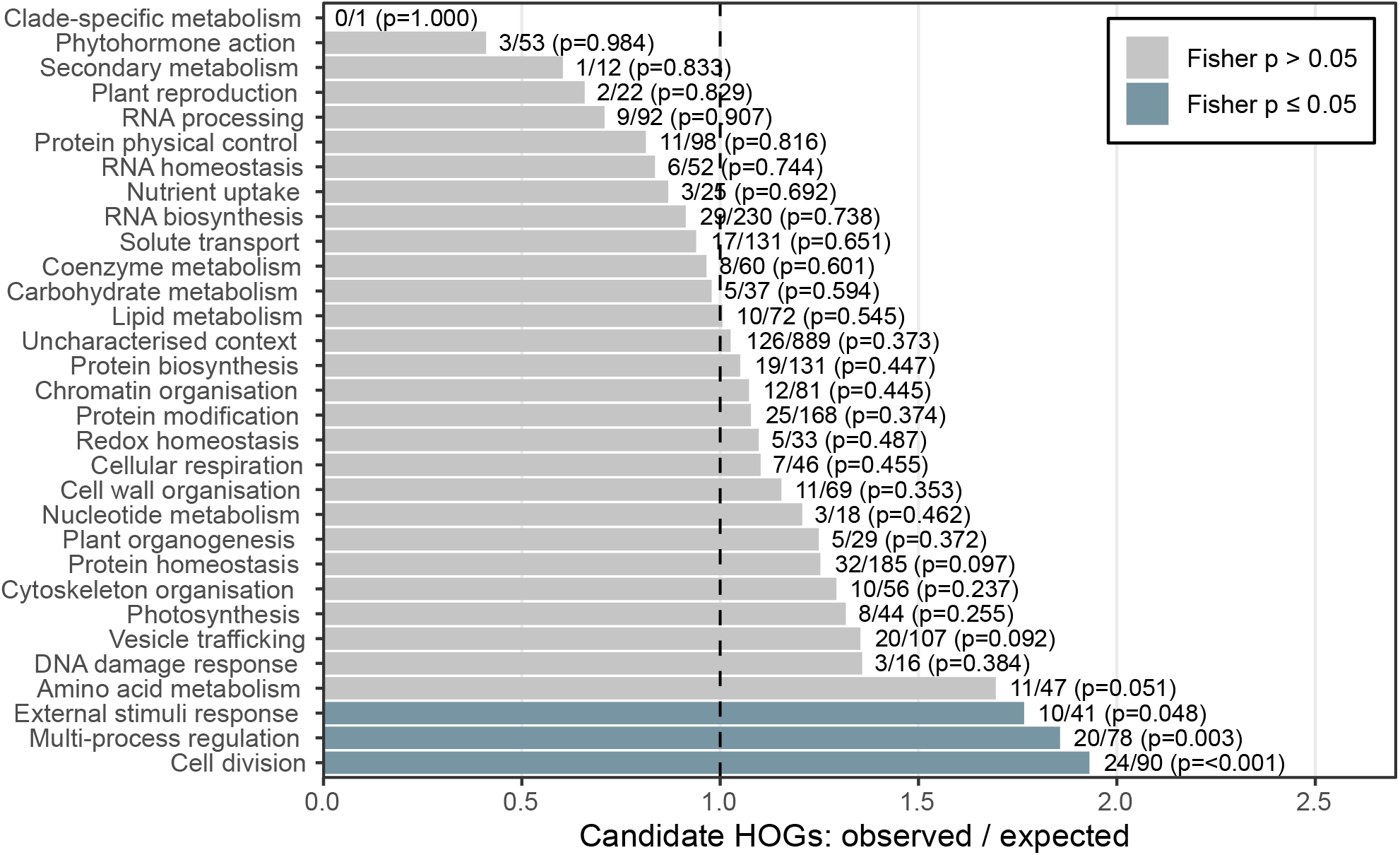
Enrichment of Mercator categories among candidate HOGs under divergent selection in annual and perennial *Hordeum* species. Bars show the observed number of candidate HOGs relative to the number expected from the background set. Labels give candidate HOGs / total HOGs in the category and the one-sided Fisher p-value.

**Figure 3:**
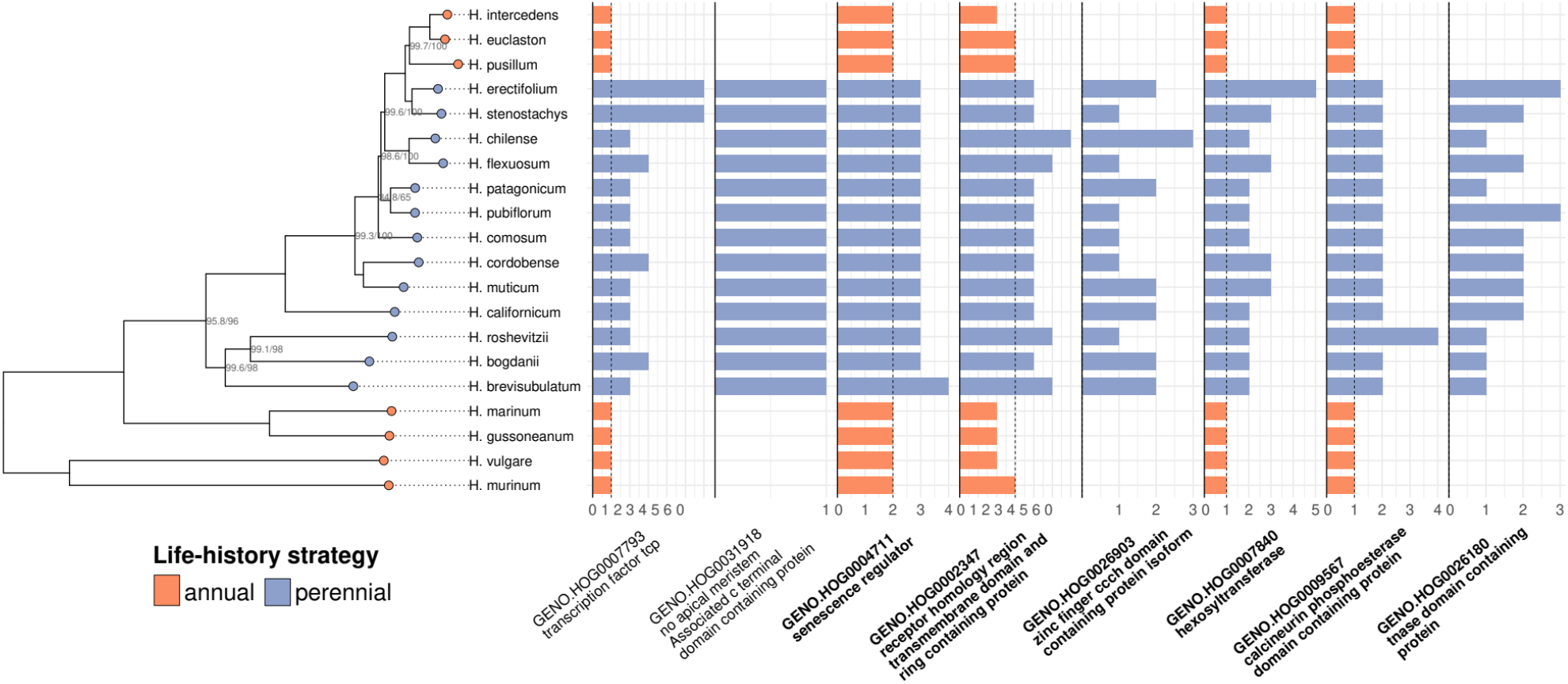
Gene copy number variation between annual and perennial *Hordeum* species. The accession tree generated from previously published genomes excluding outgroup species is shown on the left (for the complete tree, see Fig. S2). Node labels depict UFB/SH-aLRT support values and are only shown if they are not 100/100. Gene copy numbers for eight selected candidate gene families are shown on the right (for the complete list of candidate gene families, see Tab. S5). Dashed lines indicate the maximum copy number observed among annual species. Bold gene-family labels indicate families for which perennial species also expressed more paralogues than were genomically present in annual species.

The broad topology agrees well with previous reconstructions of the genus. Relationships among Old World *Hordeum* species have been consistently resolved, whereas relationships within series Critesion, particularly among the more recently diversified South American species, have remained more uncertain and were frequently characterised by polytomies in studies based on smaller sets of genetic markers (Svitashev et al., 1994; Petersen et al., 1998; Petersen and Seberg 2003; Blattner 2004; Jakob and Blattner 2006; Petersen et al., 2011; Brassac and Blattner 2015; Jin et al., 2024). Our species tree broadly corroborates these earlier topologies but differs at some positions, most notably in the placement of *H. pusillum*. Brassac and Blattner (2015) recovered *H. pusillum* as sister to *H. stenostachys* and *H. erectifolium*, a topology that would require at least one additional transition between annual and perennial life histories. Support at the nodes where the topologies disagree was substantially higher in our species tree, which contained 22,571 parsimony-informative sites compared with 2,852 in Brassac and Blattner (2015). Our topology was also congruent with the more recent species tree of Jin et al. (2024), but again showed generally higher node support than their analysis based on 7,064 parsimony-informative sites. The larger dataset and higher support herein therefore strengthen the phylogenetic basis for reconstructing life-history evolution in *Hordeum*.

Support was high throughout all three trees, with one localised exception involving *H. comosum*. Only three nodes in the Bayesian species tree had posterior probabilities below 1 (0.99, 0.94 and 0.86), all within section Stenostachys, whereas in the two accession-based ML trees the split involving H. comosum showed weaker support and represented the only topological incongruence among the three reconstructions. This instability is consistent with previous studies. *H. comosum* has alternatively been recovered as sister to *H. patagonicum* and *H. pubiflorum*, forming the youngest independent South American lineage as in our species tree, or as the second youngest independent South American lineage following divergence of the *H. muticum*-*H. cordobense* clade, as in our accession-based trees (Brassac and Blattner 2015; Jin et al., 2024; Feng et al., 2025). Increasing amounts of genomic data can strengthen support for many relationships without necessarily eliminating all phylogenetic conflict (Mason-Gamer and White 2024), and *H. comosum* appears to represent such a case in *Hordeum*. Limited within-species sampling may contribute to this uncertainty, although its cause remains unresolved. Because *H. comosum* was consistently nested within the perennial clade, its exact placement does not affect our annual-perennial comparisons performed herein.

We next tested explicitly for reticulate evolution using a two-step approach. Phylogenetic networks allowing between zero and ten reticulations identified five candidate interspecies reticulations (R1-R5; Fig. 1, S4, S5), which were subsequently evaluated using D-statistics in ABBA-BABA tests (Fig. 1; Tab. S2). Four candidates received strong support: adjusted p-values were significant in 92% of the ABBA-BABA tests for R1, 100% for R2, 89% for R3 and 100% for R4, whereas the single test of R5 was not significant. Thus, the combined network and D-statistic analyses supported four interspecies introgression events. Three of these four events were associated with major geographic disjunctions: R1 connected lineages distributed in Europe/West Asia with those in East Asia/North America, whereas R2 and R3 connected South and North American lineages (Fig. 1). This pattern suggests that long-distance dispersal may have created opportunities for hybridisation between geographically distant *Hordeum* lineages. None of the supported reticulations involved *H. comosum*, indicating that the instability of this species is not explained by the introgression events detected here. Robust inference of ancient introgression remains difficult because signals of gene flow may be restricted to particular genomic regions and can be confounded with incomplete lineage sorting (Hibbins and Hahn, 2022). We therefore restricted our interpretation to reticulations recovered across multiple network analyses and subsequently supported by D-statistics. Previous genomic studies have identified additional putative introgression events in *Hordeum*, but there is no clear agreement among studies on the identities of individual events (Jin et al., 2024; Feng et al., 2025), likely reflecting differences in genomic regions, sampling strategies and analytical frameworks. A notable exception may be our R1 reticulation, which might correspond to the deep chromosome 1H introgression recently identified by Feng et al. (2025), who found a large chromosomal segment shared between Eurasian H/Xu-genome lineages and a subset of South American I-genome species. Although differences in the used data types prevent direct assignment of the two signals to the same historical event, both analyses independently support ancient long distance gene flow between H/Xu- and I-genome lineages. Despite this uncertainty concerning individual events, the broader evidence indicates that interspecies hybridisation has occurred repeatedly during the evolution of *Hordeum*. Putative introgression has now been detected in several independent genomic studies, and numerous polyploid *Hordeum* species originated through hybridisation between distantly related lineages within the genus (Brassac and Blattner 2015; Jin et al., 2024; Feng et al., 2025). Together, these observations suggest that reticulate evolution has been a recurrent component of *Hordeum* diversification.

**Figure 4.**
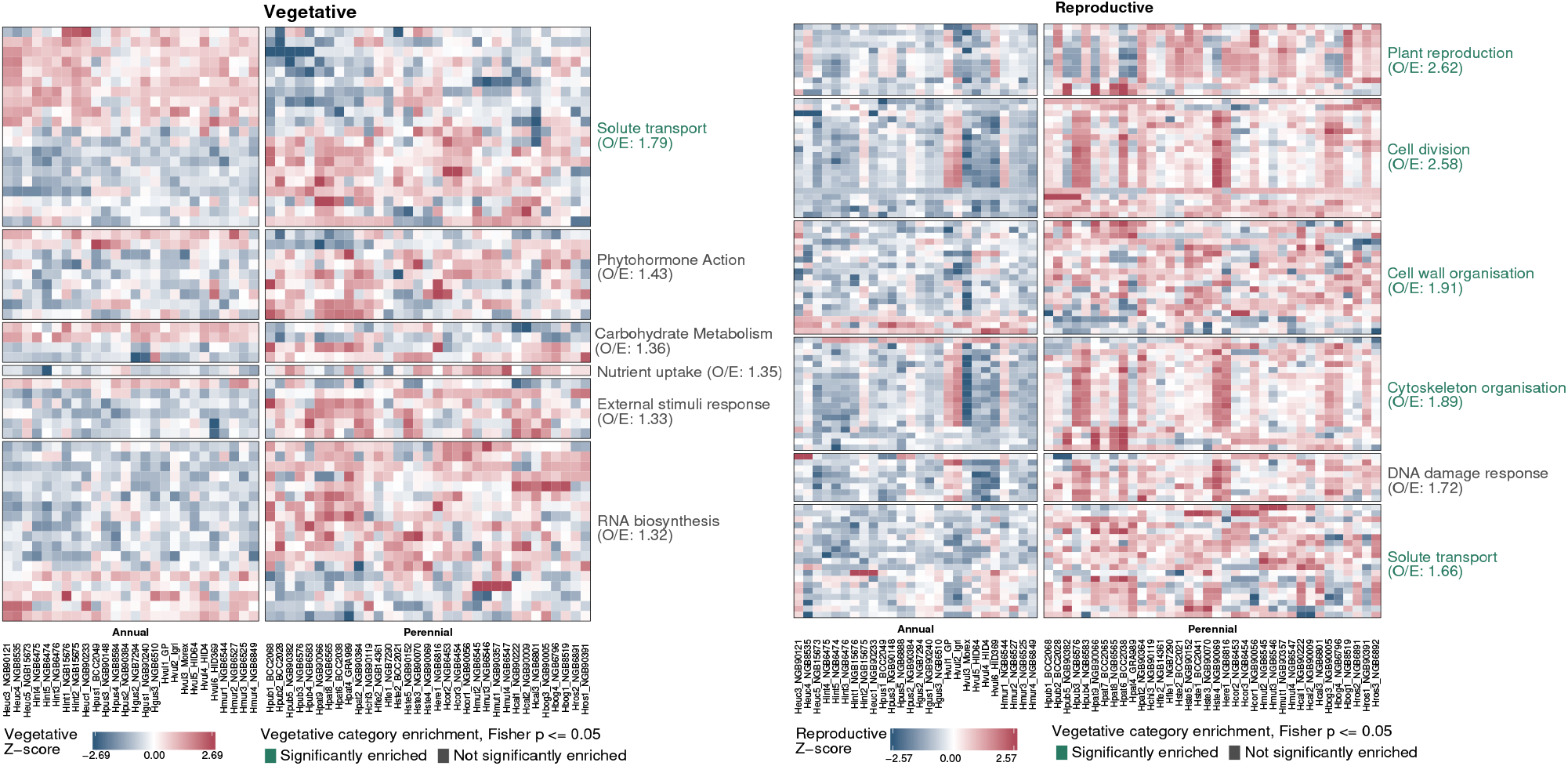
Life-history-associated gene-expression patterns in vegetative and reproductive tissues. Heatmaps show significant HOGs that are differentially expressed between annual and perennial species. Vegetative and reproductive tissues were analysed and plotted independently; consequently, the two panels contain different sets of HOGs. Columns represent accessions, separated into annual (left) and perennial (right) species and ordered within each group according to the rooted phylogenetic tree. Expression values presented as Z-scores of TMM-normalized log(2) counts per million after phylogenetic correction. Rows represent HOGs grouped by their broad Mercator functional category. Categories are ordered from highest to lowest observed/expected candidate HOG ratio, shown in parentheses after the category name. Category names are coloured when the category is significantly enriched among the differentially expressed HOGs according to a one-sided Fisher’s exact test (P =< 0.05).

**Figure 5.**
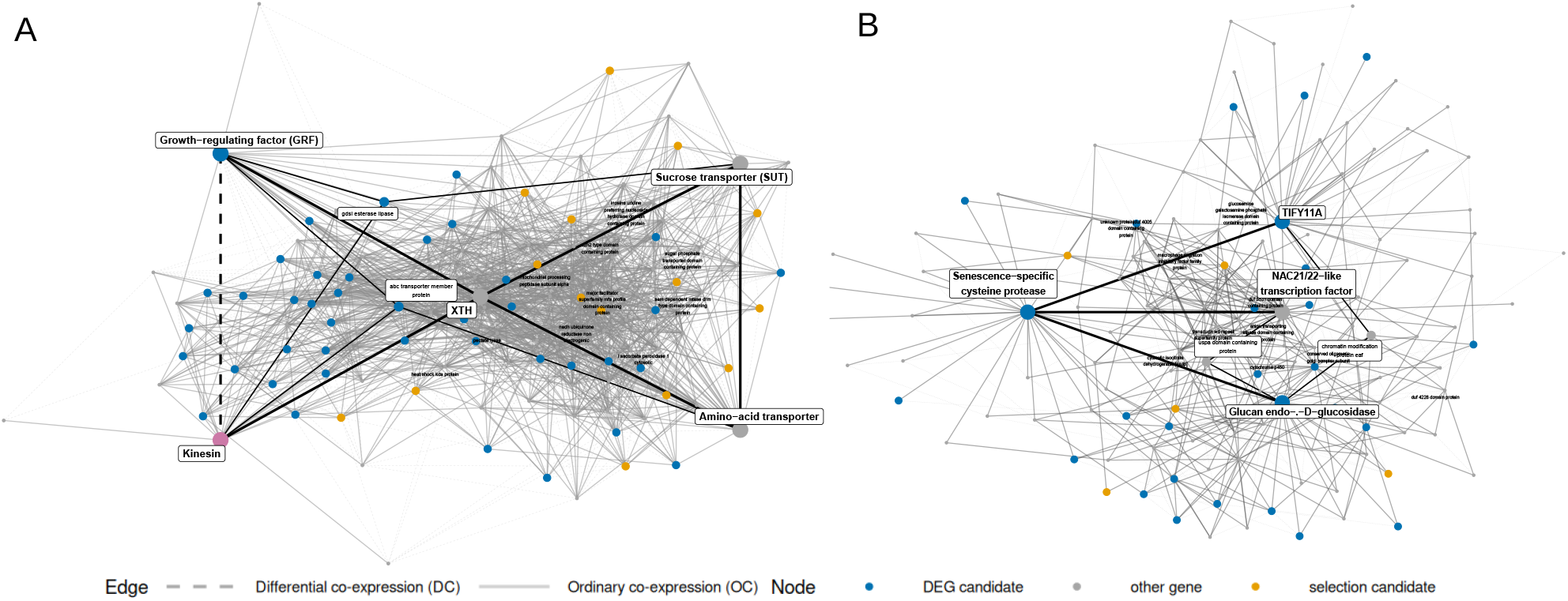
Differential co-expression networks of selected reproductive-stage genes. Boxed labels indicate the focal genes; smaller boxed labels indicate intermediate genes connecting two focal genes through a two-edge path. Black edges highlight connections among focal genes and these intermediate paths, with thicker edges used for direct focal-gene connections and thinner edges for paths through an intermediate gene. Solid edges indicate differential co-expression (DC), whereas dotted edges indicate ordinary co-expression (OC); non-highlighted connections are shown in grey. Node colours indicate differential-expression candidates, selection candidates, genes identified by both analyses, and other genes. Additional unboxed labels denote selected high-degree hubs. Node positions and edge lengths are determined by the network layout and have no quantitative meaning.

### Divergent genetic determinants in annual and perennial *Hordeum* species

Identification of genomic features distinguishing annual and perennial *Hordeum* species can help pinpoint the traits and biological processes that differ between these life-history strategies. We focused on three forms of genomic differentiation that consistently distinguish annual species in the basal and distal parts of the *Hordeum* phylogeny from perennial species, thereby separating life-history-associated patterns from effects of shared ancestry: (i) signatures of divergent selection in protein-coding sequences, (ii) gene copy number variation, and (iii) differential gene expression and co-expression networks.

#### Selection signatures

To detect signatures of divergent selection between annual and perennial *Hordeum* species, we analysed 2,908 scHOGs represented by at least two accessions per species using d_N_/d_S_ ratio modelling. After excluding nine outlier scHOGs, 573 of the remaining 2,899 scHOGs showed significant differences in d _N_/d_S_ between annual and perennial branches (BH-adjusted p ≤ 0.05; Tab. S3). In the following we focused on significant scHOGs with absolute difference between annual and perennial of d _N_/d_S_ ≤ 0.1. This resulted in 401 candidate scHOGs, with a median absolute d_N_/d_S_ difference of 0.202. Of these, 337 had higher d_N_/d_S_ in annual branches, whereas 64 had higher d_N_/d_S_ in perennial branches. For 390 of the 401 candidates, d_N_/d_S_ remained below one in both groups, indicating that the observed differences predominantly reflect changes in the strength of purifying constraint rather than pervasive positive selection. The pronounced directional asymmetry therefore suggests that protein-coding evolution was more frequently accelerated or less constrained in annual than in perennial *Hordeum* species. Importantly, genes with elevated d_N_/d_S_ in annuals showed this pattern across both independent annual lineages - the basal Old World annual lineage and the independently derived New World annual lineage - which were consistently differentiated from the perennial clade, supporting recurrent divergence in selective constraint associated with annual life history. This is broadly consistent with earlier studies reporting higher molecular evolutionary rates in annual plants (Andreasen and Baldwin 2001; Yue et al., 2010).

Mercator assigned functional categories to 399 of the 401 candidates. Three broad categories were enriched at the one-sided Fisher threshold of p ≤ 0.05: cell division, multi-process regulation, and external stimuli response (Fig. 2, Tab. S4). Cell division showed the strongest enrichment, with 24 candidates compared with 12.4 expected (1.93-fold enrichment, p = 7.8 × 10⁻⁴). These candidates included cyclins and mitotic-check - point proteins, DNA replication and repair components, kinesins and other cytokinesis-associated proteins, and proteins involved in plastid division. Their enrichment points to divergence in the cellular processes controlling proliferative growth, consistent with the different requirements for growth and tissue maintenance in annual and perennial species. Notably, several cell-division candidates directly involved in DNA replication and genome maintenance, including replication factor C, an origin-recognition-complex subunit, DNA topoisomerase, and an ATP-dependent DNA ligase, all showed higher d_N_/d_S_ in annual branches. Maintenance of meristematic stem-cell lineages is central to plant longevity (Karami et al., 2023), and remarkably low mutation rates in shoot meristems of long-lived trees suggest that these lineages are effectively protected against mutation accumulation (Schmid-Siegert et al., 2017). The higher d_N_/d_S_ of several genome-maintenance genes in annuals is therefore consistent with increased purifying constraint on genome integrity in perennials, whose cell lineages must remain functional across repeated cycles of growth and reproduction. Recent comparisons among annual and perennial Pooideae identified the maintenance of vegetative meristems beyond reproduction as a central feature distinguishing perennial growth (Hjertaas et al., 2026). Multi-process regulation contained 20 candidates (1.86-fold enrichment, p = 0.0035). This category included components of TOR (Target of Rapamycin) and SnRK1 signalling, the circadian regulator PRR1, and proteins involved in calcium, phosphoinositide, and MAP-kinase signalling, implicating regulatory systems that coordinate growth and development with metabolic processes and environmental signalling. The SnRK1-TOR system provides a well-established link between metabolic status and developmental regulation (Baena-González and Hanson 2017; Caldana et al., 2019). PRR genes in barley are core component of the circadian clock, which integrates daily and seasonal variation in light and temperature with developmental and physiological responses (Campoli et al., 2012; Ford et al., 2016; Müller et al., 2020). In perennial species, circadian-clock components also contribute to seasonal growth cessation, dormancy, cold hardiness and subsequent growth resumption (Ibáñez et al., 2010). PRR-family genes can also directly affect life-cycle strategy: in beet (Beta vulgaris), allelic variation at the pseudo-response regulator BvBTC1 determines annual versus biennial growth by modifying the response to photoperiod and vernalisation (Pin et al., 2012). External-stimulus responses contained ten candidates (1.77-fold enrichment, p = 0.048), including proteins associated with responses to light, temperature, water availability, and pathogens, such as a chloroplastic ClpB chaperone, a light-responsive bHLH transcription factor, and several protein kinases. Together, these enrichments indicate that divergent protein evolution between annual and perennial *Hordeum* species is concentrated in processes regulating cell proliferation, genome integrity and coordinating development with metabolic, environmental, and stress-related signals.

Functional enrichment highlights recurrent processes, but several candidates outside these categories encode developmental or meristem regulators, classes of genes previously linked to annual-perennial differentiation (Melzer et al., 2008; Wang et al., 2009; Zhai et al., 2024). Notable examples included a CONSTANS-like zinc-finger protein, which showed higher d_N_/d_S_ in annual branches and is associated with photoperiodic control of flowering, determining when plants transition from vegetative growth to reproduction within the growing season (Putterill et al., 1995; Campoli, Drosse, et al., 2012; Mulki and von Korff 2016). In perennial *Arabis alpina*, CONSTANS was recently shown to have distinct roles in primary and axillary shoots, which the authors hypothesise to be linked to the polycarpic life history of the species (Sashidhar & Coupland, 2026). SCARECROW, also characterised by higher d_N_/d_S_ in annuals, regulates asymmetric cell division and tissue patterning in grass meristems and may therefore be relevant to differences in the establishment and maintenance of vegetative structures between life-history strategies (Lim et al., 2000; Kamiya et al., 2003).

Two chromatin-remodelling ATPases, SYD and ISWI, likewise showed divergent d_N_/d_S_, but in opposite directions: SYD had higher d_N_/d_S_ in annuals, whereas ISWI had higher d_N_/d_S_ in perennials. These chromatin remodellers contribute to meristem identity and developmental phase transitions (Wagner and Meyerowitz 2002; Li et al., 2012), processes that must be coordinated once in annuals but repeatedly across successive growth and reproductive cycles in perennials.

Taken together, the selection results indicate that purifying constraint is generally weaker in annuals, particularly among genes involved in cell division and genome maintenance. This pattern has a straightforward life-history interpretation. In perennials, meristematic cell lineages must remain functional across repeated cycles of growth and reproduction, so mutations that compromise genome integrity can affect several future reproductive episodes and may therefore be more strongly selected against. Regulatory genes involved in developmental and environmental responses face a related constraint: perennial individuals must repeatedly adjust the same genotype to conditions that vary among years. Annuals, by contrast, complete only a single reproductive cycle, and population composition can change between years through generational turnover. Annual populations may therefore track interannual variation more rapidly (Zeineddine and Jansen 2009; Exposito-Alonso et al., 2018; Kulbaba et al., 2019). The observed excess of higher d_N_/d_S_ in annual branches is thus consistent with relaxation of purifying constraint on functions whose long-term maintenance or repeated deployment is especially important in perennials.

#### Copy number variation

Gene copy-number variation (gCNV) can affect plant phenotypes through altered gene dosage and through divergence in regulation and function following gene duplication (Ganko et al., 2007; Panchy et al., 2016). We therefore examined gCNV between life histories by testing 29,565 gene families for consistent copy-number differences between annual and perennial *Hordeum* species. We identified 41 families for which the copy-number ranges of annuals and perennials did not overlap. Although our screening allowed differences in either direction, all 41 families had higher copy numbers in perennials, with no cases of consistently higher copy number in annuals (Tab. S5).

Seven of the 41 families represented transposable-element-related proteins, including reverse transcriptases, LTR retrotransposon proteins, and transposases. Such differences are not unexpected in the repeat-rich genomes of *Hordeum* species, in which transposable elements constitute more than 80% of genomic sequence and variation in their abundance is a major contributor to genome-size differences (Feng et al., 2025). Previous comparisons within *Hordeum* also found that the relationship between genome size and life history differs among clades, indicating that life history is not linked to genome size (Jakob et al., 2004). Thus, the transposable-element-related families likely reflect broader differences in repetitive genome composition and are difficult to interpret in the context of life history strategies. Among the non-transposable-element families, several encoded proteins with potential roles in developmental regulation, including gibberellin 2-oxi-dases, laccases, flavin-containing monooxygenases, an F-box protein, and several zinc-finger proteins. Notably, Heidel et al. (2016) similarly identified gene families with corresponding predicted functions that were retained in perennial Brassicaceae but lost in annual relatives, including oxidoreductase-, F-box-, and zinc-finger-related families. Lu et al. (2026) identified 91 gene families present in three perennial *Oryza* species but absent from their annual relatives; many encoded signal-transduction-related proteins, including F-box, MYB/SANT-like proteins and kinases. Although these families are not necessarily homologous to those identified herein, the shared direction suggests that reduced representation of particular gene families may be recurrently associated with annual life history. Other *Hordeum* candidate families with higher copy number in perennials encoded developmental regulators, including two MYB transcription-factor families, a YABBY family, a no-apical-meristem-associated protein, a senescence regulator, and a TEOSINTE BRANCHED1-like TCP family. The latter is particularly interesting because TB1 regulates the dormancy and outgrowth of tiller buds in grasses; in barley, its orthologue INT-C mediates environmentally induced arrest of tiller-bud growth (Wang et al., 2022). This process is directly relevant to perenniality because, after reproductive tillers senesce following flowering, perennial plants persist through basal meristems that remain vegetative or dormant and produce replacement tillers in subsequent growth cycles (Sarath et al., 2014). In switchgrass (Panicum virgatum), progressively reducing the number of functional TB1 alleles increased axillary-bud out-growth and tiller number, demonstrating a dosage-dependent effect of TB1 on shoot architecture (Liu et al., 2020). This dosage sensitivity is particularly relevant to our result because gCNV provides a mechanism by which TB1 dosage could differ between annual and perennial *Hordeum* species.

We examined whether the additional genomic copies of perennial species were also expressed using our RNA-seq data (Tab. S5). This likely represents a conservative estimate because multimapping reads were excluded and expression was sampled across a limited set of tissues and developmental stages. For the TB1-like family, despite the higher genomic copy number in perennials, most perennial species expressed only one family member in more than half of their accessions. In six families, however, every perennial species expressed more paralogues in more than half of its accessions than the maximum number of genomic copies present in annual species. These included a senescence-regulator family, a receptor/RING-domain family, a hexosyltransferase, a calcineurin-phosphoesterase-domain protein, a transposase-domain family, and a CCCH zinc-finger family. For example, annual species contained two copies of the family annotated as a senescence regulator, whereas perennials contained three to four copies and consistently expressed three. This is particularly interesting in the annual-perennial contrast because reproduction in annuals culminates in whole-plant senescence, whereas perennial plants must restrict senescence to particular organs or shoots while maintaining meristems and other tissues for subsequent growth and reproduction (Munné-Bosch 2008; Thomas 2013). The retention and expression of additional copies of a senescence-associated gene family in perennial *Hordeum* is therefore consistent with a more complex, potentially tissue-specific regulation of senescence in perennial species. To determine whether the additional copies in perennials contribute to perennial growth in *Hordeum*, more detailed analyses are required, for example, paralogue-specific expression profiling using long-read RNA sequencing of basal buds and meristems sampled across reproductive and regrowth stages.

The consistently higher copy numbers in perennials we observed may be associated with a greater scope for regulatory or functional specialization among paralogues. This could be particularly advantageous in perennials, which must repeatedly coordinate partly contrasting developmental programmes, for example reproduction and tissue senescence with the simultaneous maintenance of vegetative meristems, followed by dormancy and renewed growth. Additional gene copies could provide the substrate for such processes to become more independently regulated across tissues or developmental stages (Ganko et al., 2007; Panchy et al., 2016).

#### Expression variation

To determine how life-history-associated gene regulation differs across development, we analysed 7,997 and 7,965 scHOGs in the vegetative- and reproductive-stage pools, respectively, using phylogenetically informed differential-expression models. The vegetative-stage pool combined roots and leaves collected from seedlings with two to three developed leaves, whereas the reproductive-stage pool combined stems and inflorescences sampled around flowering. The two pools capture distinct phases of the life cycle: early vegetative growth, when both annuals and perennials are primarily engaged in establishment and resource acquisition, and reproduction, when annuals enter a terminal reproductive cycle, whereas perennials must reproduce while retaining vegetative meristems and tissues capable of supporting subsequent growth.

Using a BH-adjusted p-value ≤ 0.05 and an absolute log_2_ fold change ≥ 1, we identified 261 differentially expressed scHOGs in the vegetative-stage pool and 618 in the reproductive-stage pool, corresponding to 3.3% and 7.8% of the tested scHOGs, respectively (Tab. S6). Of the vegetative-stage candidates, 192 were more highly expressed in perennials and 69 in annuals, whereas 540 reproductive-stage candidates were more highly expressed in perennials and 78 in annuals. Only 59 scHOGs were differentially expressed in both pools; 54 retained the same direction of expression difference and five reversed direction. Thus, most life-history-associated expression divergence was stage and tissue-specific, with a considerably stronger and more directional difference emerging during reproduction.

During vegetative growth, solute transport was the only enriched broad category, containing 20 candidates (1.79-fold enrichment, Fisher p = 0.008), equally divided between higher expression in annuals and perennials (Tab. S7). These included transporters for sugars, amino acids, nitrate, ammonium, metals, water, and other ions, indicating divergence in nutrient acquisition and resource distribution rather than a general increase in transport activity in either life history. Enriched subclasses additionally included responses to oxygen availability and ABA-related functions; both HRE/RAP2-12-like oxygen-response candidates and all three ABA-related candidates were more highly expressed in perennials. Several developmental regulators also differed individually, with an FT-like gene, a MADS-box transcription factor, SCARECROW, and a phytochrome-interacting factor more highly expressed in annuals, whereas PHYTOCHROME A and two components of the SnRK1 regulatory system were higher in perennials. SnRK1 links metabolic status to growth and developmental decisions (Baena-González & Hanson, 2017), making the higher expression of these components in perennials consistent with differences in how metabolic status is integrated into developmental regulation during vegetative growth. The recurrent involvement of SnRK1-related genes in both the expression and selection analyses suggests that annual-perennial divergence extends beyond flowering-time regulation to coordinating development with metabolic and environmental signals. This is particularly interesting given evidence from Pooideae that shifts towards greater above-ground resource allocation may precede and facilitate evolutionary transitions to annuality (Lindberg et al., 2020).

A much stronger functional pattern emerged during reproduction. Five broad categories were enriched: cell division, cell-wall organisation, cytoskeleton organisation, solute transport, and plant reproduction (Fig. 4). Cell division showed the strongest enrichment, with 52 candidates (2.58-fold enrichment, Fisher p = 1.1 × 10⁻¹⁰), followed by solute transport with 45 candidates (1.66-fold, p = 4.7 × 10⁻⁴), cell-wall organisation with 28 (1.91-fold, p = 6.4 × 10⁻⁴), cytoskeleton organisation with 26 (1.89-fold, p = 0.0011), and plant reproduction with 12 (2.62-fold, p = 0.0016). Expression within these categories was strongly biased towards perennials: 51 of 52 cell-division candidates, 24 of 26 cytoskeleton candidates, 22 of 28 cell-wall candidates, all 12 plant-reproduction candidates, and 40 of 45 solute-transport candidates were more highly expressed in perennial species.

The concurrent enrichment of cytoskeleton organisation, cell-wall organisation and solute transport places the cell-division signal in a broader developmental context. Cytoskeletal and cell-wall candidates included kinesins and formins, as well as callose synthase and pectin methylesterases, linking the expression difference to cellular processes required for cytokinesis, cell expansion and tissue formation. The cell-division category also showed the strongest enrichment among the selection candidates. Thus, the two analyses independently point to divergence in cell proliferation, at the level of both evolutionary constraint and gene activity. The strongly directional difference in expression further suggests that this divergence is particularly pronounced around reproduction, consistent with the contrasting developmental trajectories of annual and perennial species. In annuals, reproduction forms part of a terminal developmental trajectory, whereas perennials must reproduce while retaining vegetative meristems capable of sustaining subsequent growth. Tiller-level studies of perennial grasses show that reproductive tillers can senesce while vegetative tillers and axillary buds maintain the tiller population (Ott & Hartnett, 2011; Rouet et al., 2021), and recent comparisons among annual and perennial Pooideae indicate that delayed branch development contributes directly to the retention of vegetative meristems beyond flowering (Hjertaas et al., 2026). Because the reproductive-stage samples comprised stems and inflorescences rather than specifically persistent vegetative meristems, the expression data do not localise this activity to the tissues responsible for perennial regrowth. Future studies targeting basal meristems, axillary buds and newly developing tillers across reproduction and regrowth could test whether the cell-division signal observed here is specifically associated with the tissues that sustain perennial growth. Even without this tissue resolution, our data show substantially higher expression of cell-division machinery in perennial *Hordeum* around reproduction, consistent with maintained proliferative activity during this developmental phase.

Solute-transport candidates included STP- and SWEET-type sugar transporters and several AVT/LHT/AAP-type amino-acid transporters. Growing tiller tissues represent strong metabolic sinks, and coordinated carbon and nitrogen flux supports tissue production during tiller regrowth in perennial ryegrass (Schnyder & de Visser, 1999). The predominantly higher expression of these transporters in perennial *Hordeum* is therefore consistent with increased resource supply accompanying the broader proliferative signal. A PIN auxin-efflux carrier, together with several additional auxin-related candidates, further connects this pattern to developmental regulation. PIN-mediated auxin transport contributes to the control of axillary-bud outgrowth in rice (Li et al., 2022), while changes in auxin biosynthesis, transport and response are also associated with proliferative arrest of the shoot apical meristem at the end of flowering ( González-Cuadra et al., 2025), linking auxin regulation to the maintenance versus cessation of meristem activity during reproduction. Together, these enriched categories suggest that reproductive-stage divergence extends beyond core cell-cycle machinery to the structural, metabolic and regulatory processes associated with continued tissue production.

Several candidates outside the enriched categories further supported this developmental pattern, including regulators associated with meristem and shoot development. A MADS/AGL transcription factor, an SBP/SPL transcription factor, a WOX-type transcription factor and two LOF1/2-like MYB transcription factors were more highly expressed in perennials. Members of these families have established roles in meristem maintenance and pattering, and axillary-meristem formation (Yan et al., 2005; Lee et al., 2009; Lin et al., 2013; Yoshikawa et al., 2016; Tripathi et al., 2018). Because these functional assignments are based on membership in comparatively broad gene families, the specific roles of the *Hordeum* candidates remain to be established, but their expression direction is consistent with the broader enrichment of meristem maintenance, proliferative and developmental functions in perennials. A complementary signal came from genes associated with senescence. An S40-type regulator was more highly expressed in annuals, whereas three cysteine proteases were more highly expressed in perennials, arguing against a simple contrast in which annuals senesce while perennials maintain growth. This is consistent with the gCNV analysis, which showed that perennials contained a larger senescence-associated gene family and expressed more of its members. Together, these results fit the greater need in perennials to coordinate senescence with continued growth, which requires a balanced, stage- and tissue-specific senescence (Munné-Bosch, 2008; Thomas, 2013).

Taken together, the differential-expression analysis adds a developmental dimension to the genomic differences identified by the selection and gCNV analyses. During vegetative growth, annual-perennial expression divergence was comparatively limited and centred primarily on transport and environmental or metabolic regulation. Around reproduction, by contrast, the difference became substantially larger and strongly directional, with perennials showing higher expression of genes associated with cell proliferation, tissue formation and resource supply. Combined with divergent selective constraint on cell-division and genome-maintenance genes and higher copy number of several developmental and senescence-associated gene families, these results converge on a central distinction between the two life histories: annual reproduction is embedded in a terminal developmental trajectory, whereas perennial reproduction must be accommodated while preserving the cellular and developmental capacity required for subsequent growth cycles.

#### Co-expression

To determine whether annual and perennial life histories differ not only in mean gene expression but also in the coordination of expression, we compared pairwise expression correlations between annual and perennial species after accounting for phylogenetic variation. For each gene pair, the analysis tested whether the correlation across annual accessions differed from that across perennial accessions with an adjusted p-value ≤ 0.05 and an absolute difference in correlation of at least 0.5. This identified 6,135 differentially co-expressed gene pairs in the vegetative-stage pool and 167,619 in the reproductive-stage pool. Thus, divergence in co-expression increased more than 27-fold around reproduction, substantially exceeding the stage difference observed for differential expression. Overall co-expression strength differed little in the vegetative tissues, with median absolute correlations of 0.184 in annuals and 0.172 in perennials and similar proportions of strongly correlated gene pairs with |r| ≥ 0.8; 0.107% and 0.116%, respectively. In reproductive tissues, however, co-expression was markedly stronger in annuals, with a median absolute correlation of 0.250 compared to 0.190 in perennials and 0.601% versus 0.090% of gene pairs showing |r| ≥ 0.8. These results indicate that reproductive divergence extends beyond changes in mean expression to a much broader reorganisation of expression relationships.

To identify genes involved in broader network rewiring, we ranked genes by the number of differentially co-expressed partners and excluded the lower 30% of non-zero scores separately for each developmental stage, resulting in 1,816 vegetative-stage and 4,060 reproductive-stage candidate hubs (Tab. S8).

Vegetative-stage hubs were enriched for multi-process regulation (70 observed versus 54.5 expected; 1.29-fold, p = 0.0074) and solute transport (95 versus 81.2; 1.17-fold, p = 0.0366; Tab. S10). These categories directly connect the network analysis to the preceding results: multi-process regulation was also enriched among genes showing divergent selective constraint, whereas solute transport was the principal enriched category among vegetative-stage DE candidates. Thus, the same functional systems that differed in sequence evolution or mean expression also differed in how their activity was coordinated with the wider transcriptome. This suggests that annual-perennial divergence during vegetative growth involves changes not only in individual regulatory and transport components, but in the integration of signalling and resource transport into broader expression programmes.

Reproductive-stage hubs showed a broader functional pattern, with enrichment of protein biosynthesis (153 candidate hubs; 1.17-fold, p = 2.9 × 10⁻⁴), protein physical control (97; 1.19-fold, p = 0.0014), lipid metabolism (110; 1.14-fold, p = 0.0089), cellular respiration (33; 1.22-fold, p = 0.0278), and cell division (149; 1.09-fold, p = 0.0326). Within these categories, ribosome biogenesis showed the strongest subordinate enrichment, with 75 candidate hubs (1.35-fold, p = 2.7 × 10⁻⁷), while microtubule-network organisation and oxidative phosphorylation were also enriched. These processes provide complementary components of cellular growth: ribosome biogenesis supports translational capacity and plant development ( Byrne, 2009), micro-tubules are central to mitosis and cytokinesis (Rasmussen et al., 2013), and mitochondrial respiration supplies ATP and carbon skeletons required for biosynthesis and growth (O’Leary et al., 2019). Their coordinated rewiring around reproduction is therefore consistent with the life-history-associated divergence in cellular growth already indicated by the selection and differential-expression analyses. The recurrence of cell division is particularly informative: its enrichment among highly rewired genes extends the same signal to a third genomic level, showing that annual and perennial *Hordeum* differ not only in evolutionary constraint on cell-division genes and their mean expression around reproduction, but also in how proliferative machinery is coordinated with other cellular processes.

A hub gene encoding a xyloglucan endotransglucosylase/hydrolase (XTH) provides a concrete example of how this broader rewiring connects processes identified in the preceding analyses. This XTH, a member of a gene family involved in modifying cell-wall xyloglucan during growth (Hara et al., 2014), was one of the strongest reproductive-stage rewiring hubs, with 564 differentially co-expressed partners, and occupied a central position between genes associated with growth regulation and cell division on one side and solute transport on the other. It was connected to a growth-regulating factor (GRF), itself a reproductive-stage differentially expressed hub with 33 differentially co-expressed partners, and to a kinesin that was independently identified as a selection candidate and was more highly expressed in perennials during reproduction. GRF-family transcription factors regulate plant growth partly through effects on cell proliferation (Kim & Tsukaya, 2015), whereas plant kinesins have direct roles in cell division (Nishihama et al., 2002). On the other side of the network, the XTH was differentially co-expressed with prominent sucrose- and amino-acidtransporter hubs. Thus, this rewired network of differential co-expression relationships connects growth regulation, cell division and cell-wall modification with carbon and nitrogen transport - the same processes that accompanied the broader proliferative signal in the reproductive-stage differential-expression analysis. This indicates that annual and perennial life histories differ not simply in the activity of these processes, but also in how they are coordinated. Another example is a senescence-specific cysteine protease which illustrates the same phenomenon from a different functional angle. The protease was more highly expressed in perennials during reproduction and had 26 differentially co-expressed partners. These included a glucan endo-β-D-glucosidase with 52 differentially co-expressed partners, a NAC21/22-like transcription factor with 33 and a TI-FY11A protein with 25; in each case, the association with the protease was weak or negative in annuals but strongly positive in perennials. C1A cysteine proteases participate in protein degradation during barley senescence and nutrient remobilisation (Parrott et al., 2010), while jasmonate/JAZ signalling and NAC transcription factors are established components of senescence regulation (Podzimska-Sroka et al., 2015; Hu et al., 2017). The perennial-specific gains in these associations suggest tighter coordination of proteolysis, resource remobilisation and senescence-related regulation during reproduction. This supports the preceding expression and the gCNV results, arguing against a simple contrast in which annuals senesce whereas perennials suppress senescence. Instead, perennial growth appears to require more complex coordination of senescence and remobilisation with the simultaneous maintenance of tissues required for subsequent growth. In this context, the lower global co-expression observed in perennials during reproduction could be interpreted not as generally weaker transcriptional regulation, but as the result of greater compartmentalisation of the observed diverging processes.

These genes represent only a few examples of the extensive reproductive-stage network reorganisation, but illustrate how differential co-expression adds a further layer to the sequence, copy-number and expression differences identified above: life-history-associated divergence is evident not only in individual genes and functional categories, but also in the relationships among them.

## Conclusion

In our analyses, annual-perennial divergence in *Hordeum* was associated with genomic differentiation across a broad range of genes and biological functions. This breadth contrasts with the idea that transitions between life-history strategies may be initiated by comparatively few genetic changes. QTL studies have identified a limited number of major-effect loci associated with annual-perennial differences (Friedman & Willis, 2013; Gruner & Miedaner, 2021), while experimental manipulation of a few flowering and meristem regulators has shown that shifts towards perennial-like or polycarpic growth can be induced through changes in only a small number of genes (Melzer et al., 2008; Zhai et al., 2024). Our analyses address a different aspect of life-history divergence. By comparing extant annual and perennial lineages, we cannot identify the mutations that originally initiated transitions between these states. Instead, we detect genomic differences associated with established annual and perennial life histories. Relatively few genetic changes may therefore be sufficient to trigger a shift in life-history strategy, while the stable maintenance of the resulting state in natural populations involves a much broader genomic regime. The breadth of the genomic differentiation observed herein is consistent with the multidimensional phenotypic and ecological differences associated with annual and perennial growth. Across angiosperms, annuals generally develop and reproduce more rapidly and allocate more strongly to reproduction, whereas perennials tend towards more conservative resource use and persistence (Garnier, 1992; Lundgren & Des Marais, 2020; Hjertaas et al., 2023). Similar trends occur within *Hordeum*, where annual species flower earlier, accumulate more biomass early in development and thrive in different climatic areas than perennials (Anokye et al., 2026). Thus, the broad genomic differentiation identified in our analyses parallels the multidimensional ecological and phenotypic divergence between annual and perennial life histories.

Despite this functional breadth, our analyses repeatedly converged on processes associated with growth and its maintenance. Cell division was enriched among genes showing divergent selective constraint, among reproductive-stage differentially expressed genes, and among highly rewired reproductive-stage co-expression hubs. Differences in cytoskeletal and cell-wall organisation, resource transport, protein biosynthesis, and cellular respiration extended this signal to the structural and metabolic requirements of tissue production, while differences in gene copy number and expression in developmental and senescence-associated genes pointed to divergence in the regulation of growth and tissue turnover. The prominence of growth- and resource-related processes is consistent with comparative analyses across Pooideae, which suggest that shifts towards greater above-ground biomass allocation may precede and facilitate transitions to annuality (Lindberg et al., 2020). Two evolutionary scenarios are therefore conceivable: a change in life-history strategy may occur first and be followed by broader adjustment of growth and resource-allocation networks, or divergence in these traits may precede or accompany the life-history transition. Our data cannot resolve this temporal order, but they are compatible with the latter scenario. If extensive changes in growth were required only after a transition to annuality, rapid adaptive evolution of the underlying genes can be expected. Instead, most selection candidates remained under purifying selection, with the higher d_N_/d_S_ observed in annual branches predominantly reflecting weaker constraint rather than positive selection. This makes a scenario plausible in which changes in growth and resource allocation partly preceded the life-history transition, followed by further evolutionary refinement after the switch.

Irrespective of the temporal order in which individual components evolved, the broad genomic differentiation identified across our analyses is consistent with the fundamentally different developmental trajectories of annuals and perennials around reproduction.

In annual species, reproduction forms part of a terminal trajectory culminating in whole-plant senescence and death. Perennials must instead reproduce while retaining the capacity to persist through an unfavourable period and subsequently initiate another cycle of vegetative and reproductive growth. The recurrent genomic differences identified herein therefore appear to reflect this contrast between terminal reproduction and continued developmental competence. Annual-perennial divergence in *Hordeum* is thus associated not with a single genetic switch, but with broad genomic differentiation structured around whether growth terminates with reproduction or is maintained beyond it.

## Material and Methods

### Plant Material and RNA sequencing

We compiled a collection of 82 genebank accessions, which included 19 *Hordeum* and two outgroup species (Tab. S10). The seeds of these accessions were sown in freshly prepared soil mixture, 99.6 % (v/v) Mini tray soil (MIM800, Balster Einheitserdewerk, Fröndenberg, Germany) and 0.4 % (v/v) Osmocote exact standard 3-4M (Scotts Company LLC). Seeds were stratified in the dark for 14 days at 4 C° and subsequently germinated under long day conditions in walk-in phytochambers (day: 16h, 22 °C; night: 8h, 18 °C). Seedlings that developed 2-3 true leaves were moved to vernalization for 9-14 weeks (day: 8h, 4°C; night: 16h, 4°C). After vernalization, seedlings were moved back into walk-in phytochambers (day: 16h, 22 °C; night: 8h, 18 °C) and grown to maturity.

Tissue samples were taken from roots, leaves, stems and inflorescence. Root samples were taken during repotting of plants into 7×7cm pots around 14 days after transfer out of vernalization. Leaf samples were taken from the second or third fully developed true leaf at Zeitgeber (ZT) 13.5-15.5 and 21.5-23.5. Inflorescence samples were taken in the booting stage (Zadok stage 60-69) sampling only the central part of the inflorescence (middle third of the inflorescence). Stem samples were taken at the same time from the same tiller as the inflorescence sample. We removed the leaves from the stem and sampled one random internode Note that not all plants flowered within one year, and therefore, we could not sample all tissues of all plants (Tab. S10).

RNA was extracted from all tissue samples using a RNeasy kit (Qiagen, Hilden, Germany). The extracted RNA of roots and leaves were pooled, and the RNA of stem and inflorescence were pooled for each accession with equal amounts of RNA per tissue sample. Both pools per accessions were sent to Biomarker Tech nologies (BMK) GmbH (Münster, Germany) for strand-specific poly-A enriched mRNA library preparation and sequencing of min. 6Gb/sample on an Illumina Novaseq 6000 with 150bp paired-end reads.

### Transcriptome assemblies

Raw reads from the leaves-root and the stem-inflorescence pools were merged into a single fastq file per accession. After assessing the raw reads’ quality using FastQC (v0.11.9; Andrews, 2010) and MultiQC (v1.7; Ewels et al., 2016), the raw reads were filtered with fastp (v0.23.2; Chen et al., 2018; Fig. S6) using the following parameter settings: -l 50, --detect_adapter_for_pe, --cut_right, --cut_right_mean_quality 20, -c.

Transcripts were assembled denovo using rnaSPADES (v3.9.1; Bushmanova et al., 2019; Prjibelski et al., 2020) with default settings. Additionally, a genome-guided assembly was done with Trinity (v2.2.0; Grabherr et al., 2011). We ran Trinity three times with kmer 20, 26 and 32 and the parameters --SS_lib_type RF and --genome_guided_max_intron 2500. The bam files required for genome-guided assembly were created by mapping the filtered reads against the reference genomes of the respective species (Tab. S10) except for the outgroup *Psathyrostachys juncea*, which due to the lack of a reference genome, was mapped against the reference of the second outgroup *Dasypyrum villosum* (Zhang et al., 2023). Our collection contained two tetraploid species, *H. bulbosum* (autotetraploid) and *Hordeum murinum* ssp. *murinum* (allotetraploid). For autotetraploid *H. bulbosum* we arbitrarily chose genome 1 from the reference genome and treated the two accessions as diploids. Reads from *H. m.* ssp. *murinum* accessions were mapped against the allotetraploid genome. Reads that mapped to genomes 1 and 2 (herein denoted as ’G’ and ’M’, respectively, sensu Rajhathy & Morrison, 1962) were extracted and separately mapped against the separated G or M genome reference. The separated reads from the G and M genome of each *H. m.* ssp. *murinum* accessions were treated like two diploid samples for further procedure. The mapping procedure was done using STAR (v2.7.10a; Dobin et al., 2013) with default settings. In a subsequent filtering step with SAMtools (v1.3.1; Danecek et al., 2021), only uniquely mapped reads were retained.

The four assemblies per accession were merged into a single fasta file and mapped against the same reference genome used for read mapping using Minimap2 with the ‘-ax splice’ option (v2.24; Li, 2018). The longest transcript was extracted from the resulting bam file with AGAT (v0.8.0; Dainat, 2020). We used BLAST+ (v2.2.26; Camacho et al., 2009) with the parameters -max_target_seqs 1, -outfmt 6 and -evalue 1e-5 to create a database based on the concatenated proteins of the barley variety Morex (v3; Mascher et al., 2021) and *D. villosum*. This database was input for Transdecoder (v5.5.0; Haas, 2024) which we used with default parameters to predict open reading frames and collections of the longest transcript, coding and protein sequences.

### Detection and processing of orthologs

The predicted protein sequences from the 85 assemblies (77 accessions from diploid species + 2 *H. bulbosum* + 3 *H. murinum* G-genomes + 3 *H. murinum* M-genomes) were complemented with protein sequences from 13 accessions from the utilized *Hordeum* reference genomes (Feng et al., 2025). Thus, the final collection consisted of 95 accessions plus 3 ’artificial’ diploid *H. m.* ssp. *murinum* samples from in total 21 *Hordeum* and two outgroup species (Tab. S10, S11). The protein sequences of this collection were used as input for OrthoFinder (v2.5.4; Emms & Kelly, 2019) with parameters -M msa and -T fasttree. The resulting gene families will have the prefix “RNA.” in the following to distinguish them from the second OrthoFinder analysis described below.

After inspecting the resulting tree created by OrthoFinder, seven accessions did not cluster at their expected positions in the tree (marked red in Fig. S7). Note, that these seven accessions could not represent intraspecies variation as they clustered far away from the approximate position of their species in the tree. For example, *H. chilense* is a South American species, yet two of its samples clustered deep in the clade of an Asian species (*H. roshevitzii*). We consider it to be likely, that all of these seven plants did not belong to the taxa they were assigned to and, therefore, represented faulty phylogenetic signals. Since these accessions may have biased the detection of orthologs, we removed them and repeated the analysis with the pruned set of 87 accessions plus three ’artificial’ diploid *H. m.* ssp. *murinum* and continued with this pruned set of accessions.

After running OrthoFinder again with the pruned set of accessions, we extracted single-copy hierarchical orthogroups (scHOGs) which were present in all accessions (257 scHOGs) and aligned their coding sequences to create a multiple sequence alignment (MSA) for each scHOG. Alignment was done with MAFFT (v7.407; Katoh et al., 2002). Each MSA was trimmed using Clipkit (v1.4.1; Steenwyk et al., 2020) with default parameters to increase the signal-to-noise ratio and improve the MSAs’ quality for subsequent phylogenetic analyses.

### Phylogenetic tree inference

We created partition schemes from the 257 scHOGs present in all accessions using PartitionFinder (v2.1.1; Lanfear et al., 2017) using raxml, a greedy search across all models and aicc as a decision criterion for model selection.

Fasta files were created that contained the scHOG sequences from the determined best fitting partitioning scheme and partition-trees of each partition were created using IQtree (v2.2.0; Minh et al., 2020) with 1000 ultrafast bootstrap values (UFB). Substitution models were determined with ModelFinder (vNA; Kalyaanamoorthy et al., 2017). The partition-trees and the MSAs were used to estimate a number of properties and sorting the partitions according to their usefulness for phylogenetic inferences following the method of the R package genesortR (Koch 2021).

None of the partitions exhibited any properties that suggested they may introduce a bias (Fig. S8; Tab. S13). Hence, all partitions were concatenated and used to create an accession-based phylogenetic tree using the maximum-likelihood approach implemented in IQtree (v2.2.0; Minh et al., 2020) with the best fitting models of nucleotide substitution determined by ModelFinder (vNA; Kalyaanamoorthy et al., 2017). We used 100,000 UFB and 100,000 bootstrap replicates of the SH-like approximate likelihood ratio test (SH-aLRT; Guindon et al., 2010) to test the robustness of the inference. Additional applied parameters were -m MFP+ and --sampling GENESITE.

A Bayesian analysis with parallel inference of the species tree and divergence times is computationally expensive and was not feasible with all scHOGs. Therefore, we only used coding sequences of the first 50 partitions (= 90 scHOGs), according to the order determined by the genesortR analysis, for Bayesian tree inference with the StarBeast3 model (v1.1.8; Douglas et al., 2022) of BEAST (v2.7.5; Bouckaert et al., 2019). For the analysis with BEAST, we implemented the substitution models determined to best fit to the 50 partitions by PartitionFinder in the above described analysis. As clock model of the species tree, we used a relaxed clock and estimated its standard deviation and clock rate. As a model of speciation, the calibrated Yule model was used. Estimations of mutation rates in wild *Arabidopsis thaliana* populations were reported to range between 2 and 5 x10^-9^ site^-1^year^-1^ (Exposito-Alonso et al., 2018). Even though it is likely that the mutation rates in perennial *Hordeum* species are slightly lower, these estimates are a sufficient proxy for a prior distribution. Therefore, we defined the priors for the clock rates of the species tree and the partitions as a log-normal distribution with a mean (µ) of 0.0035 and a standard deviation of 0.9 (note, these values were transformed to site^-1^ [million years]^-1^). We created three MRCA priors for time calibration. A normal distribution with µ=19.4 and =6 at the split of *P. juncea* and *D. villosum* (Bernhardt et al., 2017), a normal distribution with µ=12 and =3 at the split of *D. villosum* and *Hordeum* (Brassac and Blattner 2015; Bernhardt et al., 2017) and a normal distribution with µ=9.23 and =1.3 at the *Hordeum* root (Bernhardt et al., 2017; Brassac & Blattner, 2015). Chain lengths were 700 million, storing every 4,000 samples. We used CoupledMCMC (1.2.0; Müller & Bouckaert, 2020) with four chains. All other parameters were left at default. The corresponding xml file can be found in the data repository. We ran this analysis three times with different seeds and inspected the log files in Tracer (v1.7.2; Rambaut et al., 2018) to ensure convergence. The three independent runs were combined with Logcombiner (v2.7.6; Rambaut et al., 2018) discarding 10% of the trees of each run as burn-in to create the final collection of tree samples. Treeannotator (v2.7.6; Drummond & Rambaut, 2007) was used to summarize tree samples to a maximum clade credibility tree with 0% burn-in and 0.5 posterior probability limit.

### Interspecies hybridization

We extracted all coding sequences from scHOGs in *Hordeum* which were present in at least two accessions per species or in one accession for the species which were represented by only a single accession. This resulted in a set of 2,909 scHOGs, which in the following will be referred to as the ‘M2A-set’. The sequences were aligned and trimmed using the same approach outlined above with the scHOGs present in all accessions.

Phylogenetic network inference was conducted with the maximum pseudo-likelihood approach developed for SNAQ (Solís-Lemus et al., 2017) and implemented in the Julia (v1.10.2; Bezanson et al., 2017) package PhyloNetworks (v0.16.3; Solís-Lemus et al., 2017). Genetrees from the aligned coding sequences of the M2A-set were estimated with IQtree (v2.2.0; Minh et al., 2020) using the same procedure as outlined above for creating the input of genesortR (Koch 2021). Species quartet concordance factors were calculated from these genetrees. We ran SNAQ ten independent times with different seeds, allowed reticulations ranging from 0-10 and using the species tree as a guide tree. We considered all networks with a pseudo log-likeli - hood below a certain threshold as candidate networks. This threshold was arbitrary. Yet, in all models, there was a steep increase in pseudo-log-likelihood between networks below and above this threshold, suggesting a clear decrease in statistical support for networks above the threshold (Fig. S4). Reticulations that appeared in at least two independent networks were considered candidate reticulations (Fig. S5).

For SNP calling, we used the reference genome from the species that was on average the least diverged from all other species. The most suitable reference genome was determined by calculating patristic distances between all accessions from the accessions tree. The species with the lowest average patristic distances to all other species was *H. comosum*, which therefore was used as the common reference genome. Filtered reads from all accessions subjected to RNA sequencing were mapped against the *H. comosum* reference genome (Feng et al., 2025) using STAR (v2.7.10a; Dobin et al., 2013) with default settings. In subsequent filtering steps with SAMtools (v1.3.1; Danecek et al., 2021), only uniquely mapped reads were retained. Variant calling was done with BCFtools (v1.10.2; Danecek et al., 2021) and default parameters. We kept only biallelic SNPs without missing data, QUAL ≥ 30, GQ ≥30, DP ≥ 5 and for heterozygous SNP calls DP ≥ 30. VCFtools (v0.1.16; Danecek et al., 2011) and BCFtools (v1.10.2; Danecek et al., 2021) were used for variant filtering. The 13 included accessions from Feng et al. (2025) were included by extracting the gene space defined in the gff files of the corresponding species from the respective reference genomes. These sequences were mapped against the *H. comosum* reference genome using Minimap2 with the asm20 option to account for the divergence between species (Li 2018). Only sequences with MAPQ ≥ 30 were used to call biallelic, homozygous SNPs with QUAL ≥ 30. The two VCF files were merged and the merged file was used for the subsequent procedure.

Linkage disequilibrium (LD) pruning was done with PLINK (v 1.90b7; Purcell et al., 2007). Windows of 50 kb were shifted by five variants, and SNPs with r^2^ ≥ 0.5 within one window were removed. The pruned SNP set was used as input for Dsuite (v0.5r53; Malinsky et al., 2021) to conduct ABBA-BABA tests for candidate reticulations found with SNAQ (Solís-Lemus et al., 2017). In all tests, *Dasypyrum villosum* was the outgroup species. Because each ABBA-BABA test requires a specific four-lineage topology, candidate reticulations could be tested with multiple species combinations when several species satisfied the required phylogenetic relationships. Consequently, the number of ABBA-BABA tests differed among candidate reticulations depending on the number of valid species combinations. In Dsuite, we used the --ABBAclustering parameter to test whether ABBA patterns associated with significant D values tend to cluster together (Koppetsch et al., 2024).

### Selection signatures in the context of life-history strategies

Coding sequences from the M2A-set were subjected to codon-aligned sequence alignment using MACSE (v2.07; Ranwez et al., 2018) with parameter -local_realign_init 1 and -local_realign_dec 1. Each of these sequences were used as input to CODEML implemented in PAML (v4.10.7; Yang, 2007) to model d_N_/d_S_ across the accession tree, including only accessions from Hordeum species, which allowed us to control for phylogenetic non-independence. We ran the null model (M_0_) and the branch model (M_alt_), in which accessions from annual species form the foreground branches.

Differences between M_0_ and M_alt_ were tested using likelihood-ratio tests, and the resulting p-values were corrected for multiple testing with the Benjamini-Hochberg method (Benjamini and Hochberg, 1995). HOGs with missing or non-finite foreground or background d_N_/d_S_ estimates, negative estimates, or estimates > 5 were excluded as we assumed these unreliable. HOGs with an adjusted p-value ≤ 0.05 were initially considered candidates. To exclude statistically significant differences with only small changes in d_N_/d_S_, we calculated the absolute difference between the background and foreground d_N_/d_S_ estimates and removed significant HOGs with that difference was smaller than 0.1. The remaining HOGs were considered candidate genes showing divergent selective constraint between annual and perennial *Hordeum* species.

Functional annotations for the analysed HOGs were based on the corresponding proteins of the *H. gussoneanum* accession BCC2005 (Feng et al., 2025). Protein sequences were submitted to Mercator with Swiss-Prot annotations enabled, and the resulting MapMan (v4; Schwacke et al., 2019) annotations were assigned to the respective HOGs. Functional enrichment was tested separately for broad MapMan classes and subclasses using one-sided Fisher’s exact tests. The enrichment background comprised all HOGs retained after d_N_/d_S_ quality filtering and represented in the Mercator output, whereas the candidate set comprised the retained candidate HOGs defined above. Each HOG was counted only once within a functional category. Categories were considered enriched at p ≤ 0.05. Expected candidate numbers and fold enrichment were calculated from the proportion of candidates in the complete enrichment background and were used as descriptive measures of enrichment.

### Gene copy number variation in the context of life-history strategies

For analyses of gene family sizes, we only used protein sequences from all reference genomes published by Feng et al. (2025) and added six outgroup species: *Arabidopsis thaliana*, *Oryza sativa*, *Zea mays*, S*orghum bicolor*, *Brachypodium distachyon* and *Dasypyrum villosum* (Tab. S12). The protein sequences of this collection were used as input for OrthoFinder (v2.5.4; Emms & Kelly, 2019) and processed the same way as out - lined above for the protein sequences from the transcriptome assemblies and the selected accessions from Feng et al. (2025). The resulting gene families will have the prefix “GENO.” to distinguish them from the first OrthoFinder analysis described above.

Single copy HOGs were extracted from the output and used for the same phylogenetic analyses like described above for the protein sequences from the transcriptome assemblies and the selected accessions from Feng et al. (2025) except for the species tree inference, i.e. alignment and trimming ➝ creation of partition ➝ estimating gene properties for phylogenetic inference ➝ accession based phylogenetic inference. The resulting accession tree is called ‘genome accession tree’.

The outgroups and all polyploid species were removed from the assignment of proteins to gene family done by OrthoFinder (v2.5.4; Emms & Kelly, 2019). We then counted the number of genes of the remaining diploid *Hordeum* species and filtered this data set. The data set contained seven annual *Hordeum* species, and we wanted to reduce the computational expenses by excluding gene families with large variations of presence/absence within annuals or perennials because these were unlikely to be functionally linked to differences between life history strategies. We allowed 1-species error margin and, hence, retained only gene families where at least 6 species had a gene count of ≥ 1.

A gene family was considered a candidate if the gene count of the smallest family in an annual species exceeded the gene count of the largest family in a perennial species or vice versa. The procedure was done in R (v4.3.3; R Core Team, 2024) using base functions. Gene features of these final candidate genes were extracted from the *H. gussoneanum* accession BCC2005 (Feng et al., 2025) for consistency with expression and selection analyses. Gene features were extracted from one representative gene per family.

### Differential gene expression in the context of life-history strategies

For the analysis of differential gene expression, only diploid *Hordeum* accessions subjected to RNA sequencing were used. Reads from the vegetative pool, consisting of leaf and root RNA, and the reproductive pool, consisting of stem and inflorescence RNA, were analysed separately. Reads were mapped against the reference genome of the respective species using STAR (v2.7.10a; Dobin et al., 2013), retaining only uniquely mapped reads. Gene-level read counts were obtained with HTSeq (v2.0; Putri et al., 2022) using reverse-stranded counting. Gene counts were subsequently assigned to HOGs based on the OrthoFinder analysis described above in the gCNV paragraph. To ensure that expression was compared between single orthologues across species, only HOGs represented by one gene in each of the reference genomes used for the expression analysis were retained.

Differential expression between annual and perennial accessions was analysed separately for the vegetative and reproductive pools using edgeR (v4.2.2; Chen et al., 2025). To account for phylogenetic non-independence among accessions, cophenetic distances were calculated from the accession-based ML tree and subjected to principal coordinate analysis with Lingoes correction using the R package ape (Paradis & Schliep, 2019). The first three principal coordinate axes were included as covariates together with life-history strategy in the statistical model. HOGs with low read counts were removed separately for each tissue pool using filterByExpr with a minimum count of 10, a minimum total count of 15 and a minimum proportion of 0.7. Library sizes were normalised using the trimmed mean of M-values (TMM) method. Dispersions were estimated using robust estimation, and quasi-likelihood negative-binomial generalized linear models were fitted with glmQLFit. Differential expression between annual and perennial accessions was tested using quasi-like-lihood F-tests. P-values were corrected for multiple testing with the Benjamini-Hochberg method (Benjamini & Hochberg, 1995), and HOGs were considered differentially expressed when the adjusted p-value was ≤ 0.05 and the absolute log_2_ fold change was ≥ 1.

Similar to the approach used in the selection analysis, functional annotations were based on the corresponding proteins of the *H. gussoneanum* accession BCC2005 (Mascher et al., 2025). Protein sequences representing the union of HOGs tested in the vegetative and reproductive analyses were submitted to Mercator4 using Swiss-Prot annotations, and the resulting MapMan4 functional assignments were linked to the respective HOGs (Schwacke et al., 2019). Functional enrichment was performed separately for the vegetative and reproductive candidate sets at both broad MapMan class and subclass levels. The background for each analysis consisted of all HOGs tested in the respective differential-expression analysis and represented in the Mercator output. Each HOG was counted only once per functional category, and enrichment was tested using one-sided Fisher’s exact tests. Categories with p ≤ 0.05 were considered enriched.

### Differential gene co-expression in the context of life-history strategies

To test whether annual and perennial *Hordeum* species differed in the coordination of gene expression, we performed differential co-expression analysis separately for the vegetative and reproductive tissue pools. HOGs with insufficient read counts were filtered separately for both tissue pools using the same criteria as in the differential-expression analysis, and only HOGs passing the filter in both pools were retained. Read counts were TMM-normalised using edgeR (v4.2.2; Chen et al., 2025) and transformed to log_2_ counts per million with a prior count of one. To account for phylogenetic non-independence among accessions, expression variation associated with the first three principal coordinate axes of the cophenetic distances from the accession-based ML tree was removed using removeBatchEffect implemented in limma (Ritchie et al., 2015), while retaining the mean expression difference between annual and perennial accessions.

Because testing all pairwise correlations among the retained HOGs was computationally expensive, the differential co-expression analysis was restricted to 6,000 HOGs. All HOGs identified as differential-expression or selection candidates and passing the expression filter were retained. The remaining positions were filled with the HOGs showing the highest expression variance, ranked according to their mean variance across the phylogenetically adjusted vegetative and reproductive expression matrices. HOGs without expression variation in either tissue pool were removed. Differential co-expression between annual and perennial accessions was then analysed separately for each tissue pool using the R package DGCA (McKenzie et al., 2016). Pearson correlations were calculated for all HOG pairs separately across annual and perennial accessions and compared between the two groups. P-values for differences in correlation were corrected for multiple testing using the Benjamini-Hochberg method (Benjamini & Hochberg, 1995). Gene pairs were considered differentially co-expressed when the adjusted p-value was < 0.05 and the absolute difference between the perennial and annual correlation coefficients was ≥ 0.5.

To identify genes involved in broader co-expression rewiring, we counted the number of differentially co-expressed partners for each HOG separately within the vegetative and reproductive analyses. HOGs without differentially co-expressed partners were excluded from the hub ranking, and the 30% of HOGs with the lowest non-zero numbers of differentially co-expressed partners were removed. The remaining HOGs were considered candidate co-expression hubs. Functional annotations were assigned using the Mercator4/Map-Man4 framework described above (Schwacke et al., 2019). Functional enrichment of candidate hubs was tested separately for vegetative and reproductive tissues at the broad MapMan class and subclass levels using one-sided Fisher’s exact tests. The background consisted of all HOGs included in the differential co-expression analysis and represented in the Mercator output. Each HOG was counted only once within a functional category, and categories with p ≤ 0.05 were considered enriched.

## Data and resource availability

Raw data, a selection of intermediate results, supplements and scripts were stored in the DataPlant Hub https://git.nfdi4plants.org/timo.hellwig/evolution_and_divergent_geneticmechanisms_of_annual_and_perennial_hordeum_species. [will be updated upon acceptance]

## Acknowledgements

We gratefully acknowledge computational support from the Zentrum für Informations-und Medientechnologie (ZIM), in particular the HPC team operating the HILBERT high-performance computing cluster at Heinrich-Heine-Universität Düsseldorf.

FAIR data management was facilitated by the DataPLANT infrastructure (Weil et al., 2023), including its personal assistance network and integrated tool stack. We particularly thank CEPLAS data steward Dominik Brillhaus for his support in creating, structuring, and curating the ARC repository associated with this study, thereby ensuring the FAIRness and long-term reusability of the underlying research data and metadata.

This work was funded by the Deutsche Forschungsgemeinschaft (DFG) under (1) Germany’s Excellence Strategy (EXC-2048/1, Project ID: 390686111) and (2) the Collaborative Research Centre/Transregio (TRR341, Project ID: 456082119).

## Authors’ Contributions

TH: Conceptualization, Investigation, Methodology, Software, Formal analysis, Visualization, Writing - original draft, Data curation, Project administration. ND: Investigation. EBH: Methodology. MvK: Conceptualization, Methodology, Funding acquisition, Resources, Writing - review & editing, Project administration, Supervision.

## Supplementary Figures

Figure S1: Accession based maximum likelihood tree. Node labels depict UFB/SH-aLRT values. Node labels are only shown if they were not the maximum of 100/100.

Figure S2: Maximum likelihood phylogenetic tree based on previously published genomes. Toemphasize the branching order despite highly variable branch - lengths, the tree is rendered as a cladogram.

Figure S3: Species tree with mean (A) and 95% HPD interval (B) of divergence times as node labels.

Figure S4: Pseudo-log-likelihood of phylogenetic network analysis using SNAQ averaged over repetitions (A) and per per number of reticulations (B-G) with networks below the threshold (green dotted line). Selected candidate networks are depicted in Fig. S5.

Figure S5: Candidate phylogenetic networks (A-H). Coloured lines indicate major (dark blue; γ > 0.5) and minor parent edges (light blue; γ < 0.5) of putative hybridization events. Their corresponding numbers represent the inheritance proportion (γ).

Figure S6: Summary statistics of the read filtering process. Each dot represents one accession.

Figure S7: Phylogenetic tree of the unpruned set of accessions created by fasttree during the first OrthoFinder analysis. Accessions in red were removed due to their possible mislabeling.

Figure S8: Value of the six gene-partition properties against their sorted position. Order was determined according to their phylogenetic usefulness estimated by genesortR.

## Supplementary Tables

Table S1: Summary statistics of single copy hierarchical orthogroups used for inference of three phylogenetic trees.

Table S2: ABBA-BABA test results of selected candidate reticulations

Table S3: List of orthogroups with output statistics of the selection analysis.

Table S4: Mapman enrichement statistics of the selection analysis seperated by Mapman class /subclass and tissue.

Table S5: Gene copy number of each species and description of one representative *H. gussoneanum* gene of each gene family that showed copy number variation between annual (red) and perennial (blue) *Hordeum* species.

Table S6: Output statistics of the differential expression analysis separated by tissue pool.

Table S7: Mapman enrichement statistics of the differentially expression analysis seperated by Mapman class /subclass and tissue.

Table S8: Output statistics of the differential co-expression analysis separated by tissue pool.

Table S9: Mapman enrichement statistics of the differentially co-expression analysis seperated by Mapman class /subclass and tissue.

Table S10: List of plant material used for transcriptome assemblies.

Table S11: Pan-Hordeum consortion (Feng et al., 2025) accessions used to complement the detection of orthologs

Table S12: Data sources of reference genomes used for creating the maximum likelihood phylogenetic tree based on previously published genomes(Fig. 3, S2)

Table S13: Summary statistics of scHOG partitions inferred by genesortR.

## Notes

### Competing Interest Statement

The authors have declared no competing interest.

### Summary of Updates

The manuscript has been substantially revised: the climatic niche analysis was removed; the genomic analyses were expanded and restructured, including separate analyses of vegetative and reproductive gene expression and the addition of differential co-expression analysis; the selection and gene copy-number analyses, interpretation, figures, and supplementary materials were revised accordingly.

