## Supplementary_Figures for "Comparative phylogenomics reveals multilevel genomic differentiation between annual and perennial *Hordeum* species"

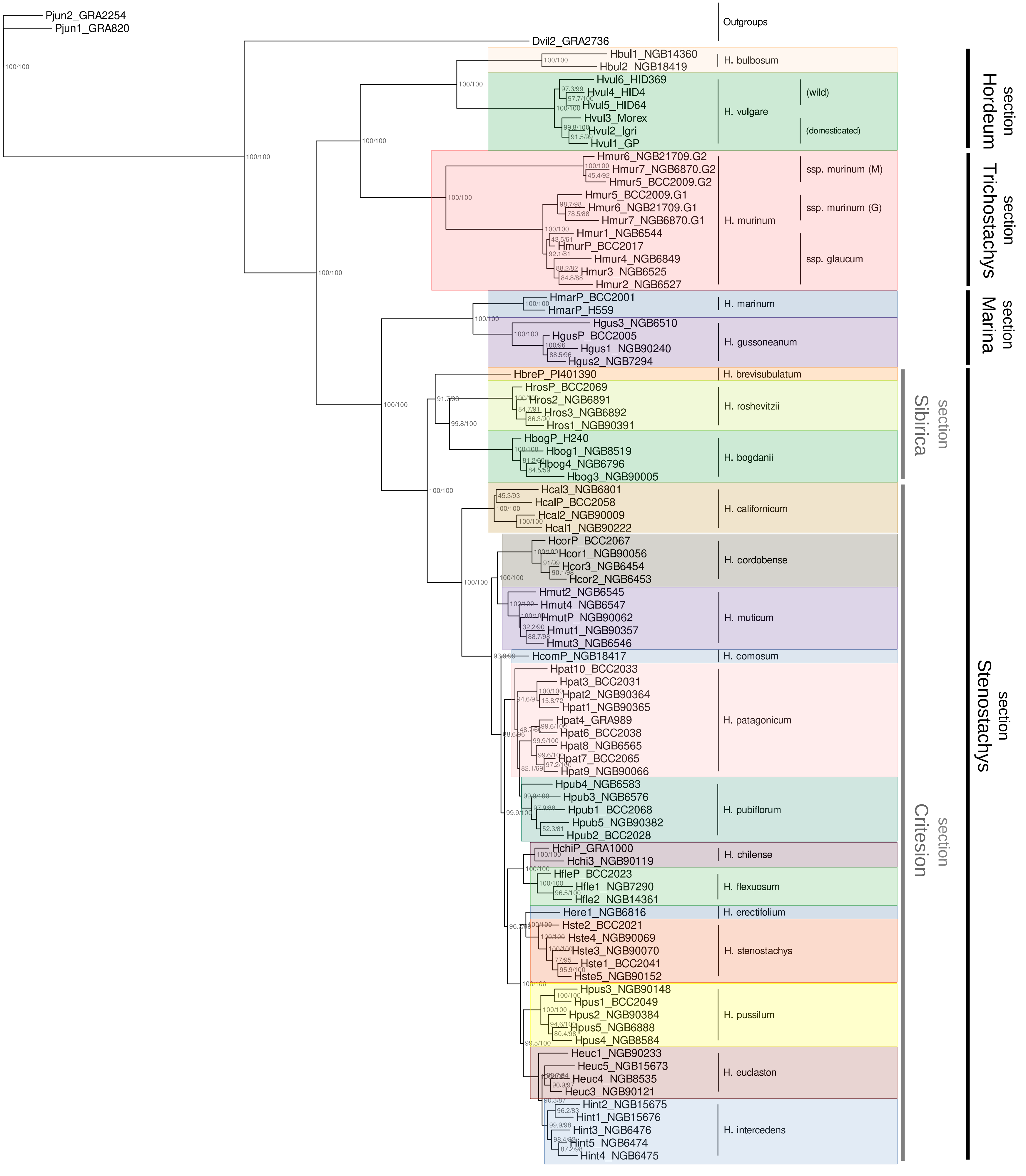


**Figure S1:** Accession based maximum likelihood tree. Node labels depict UFB/SH-aLRT values. Node labels are only shown if they were not the maximum of 100/100.

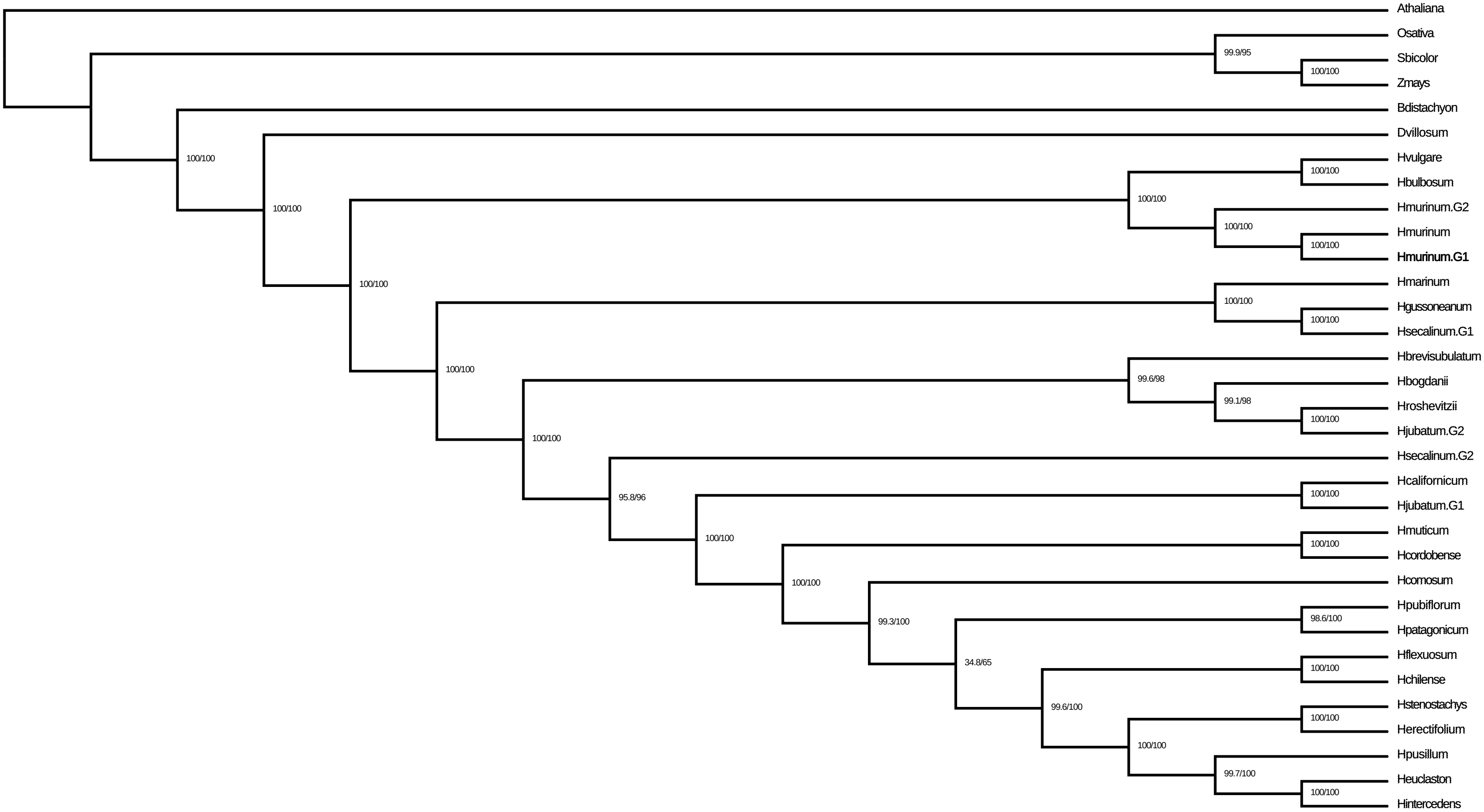


**Figure S2:** Maximum likelihood phylogenetic tree based on previously published genomes. To emphasize the branching order despite highly variable branch-lengths, the tree is rendered as a cladogram.

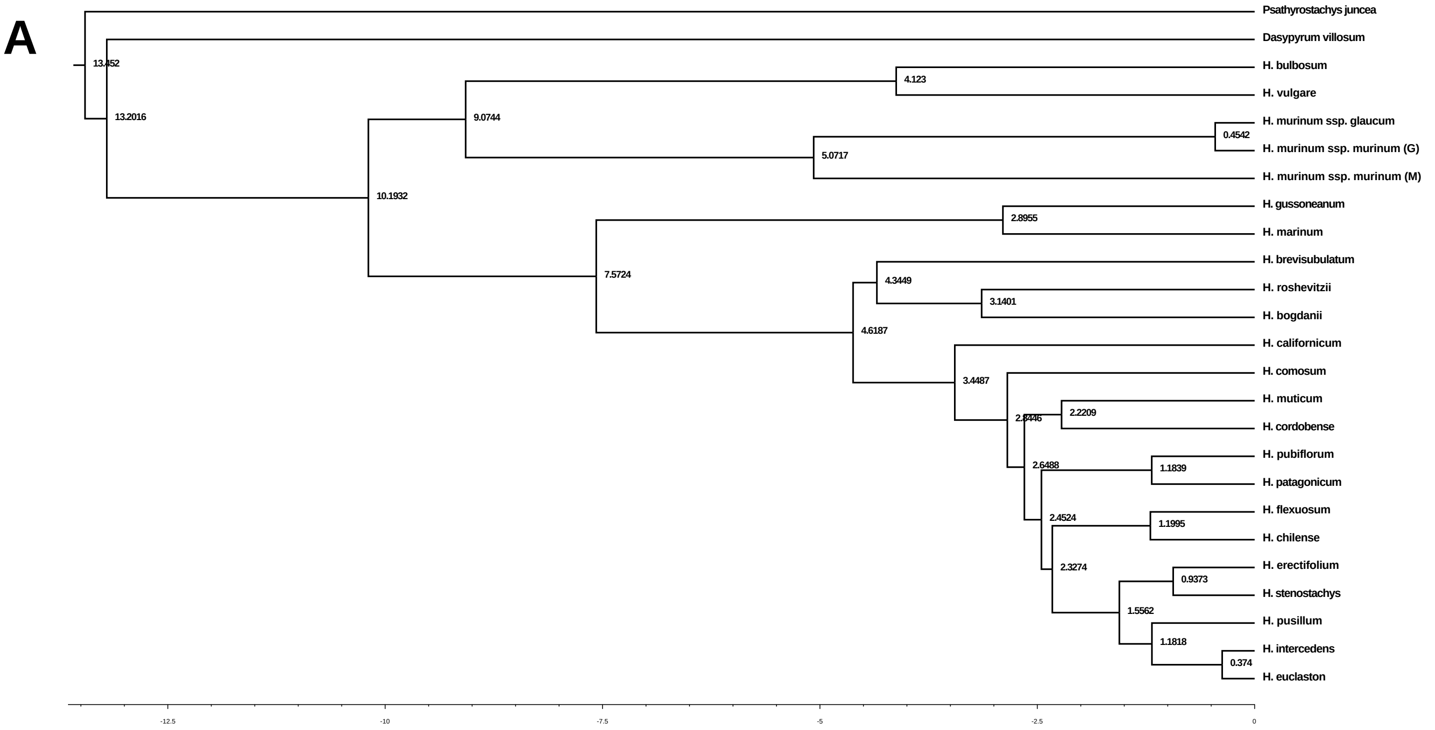

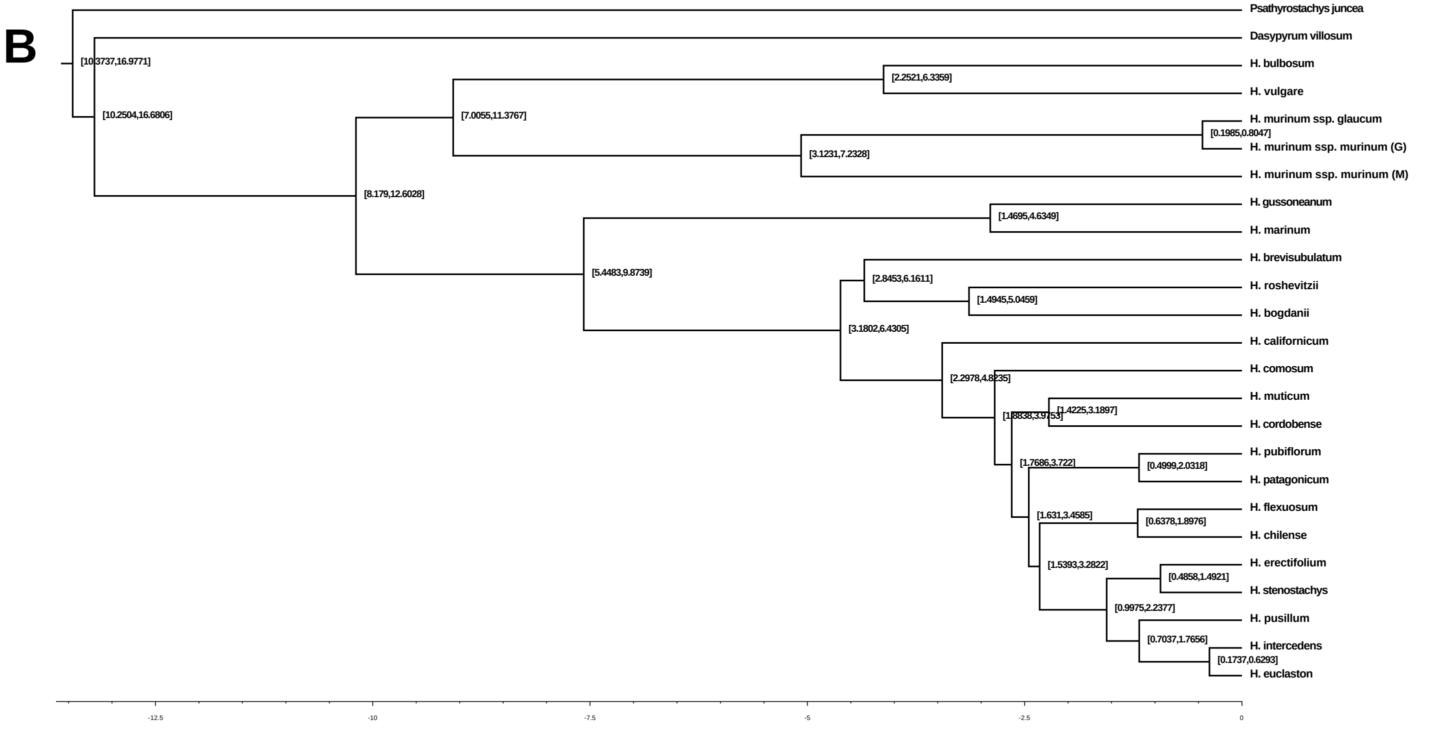


**Figure S3:** Species tree with mean (A) and 95% HPD interval (B) of divergence times as node labels.

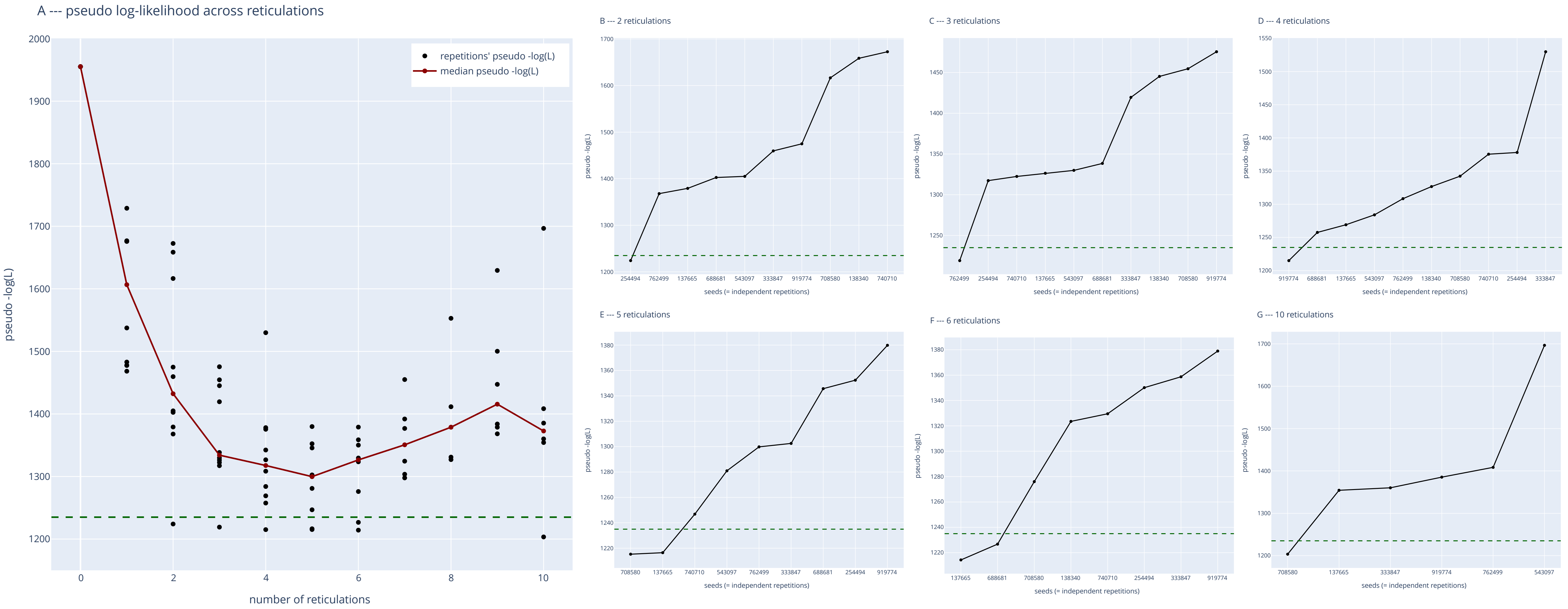


**Figure S4:** Pseudo-log-likelihood of phylogenetic network analysis using SNAQ averaged over repetitions (A) and per per number of reticulations (B-G) with networks below the threshold (green dotted line). Selected candidate networks are depicted in Fig. S5.

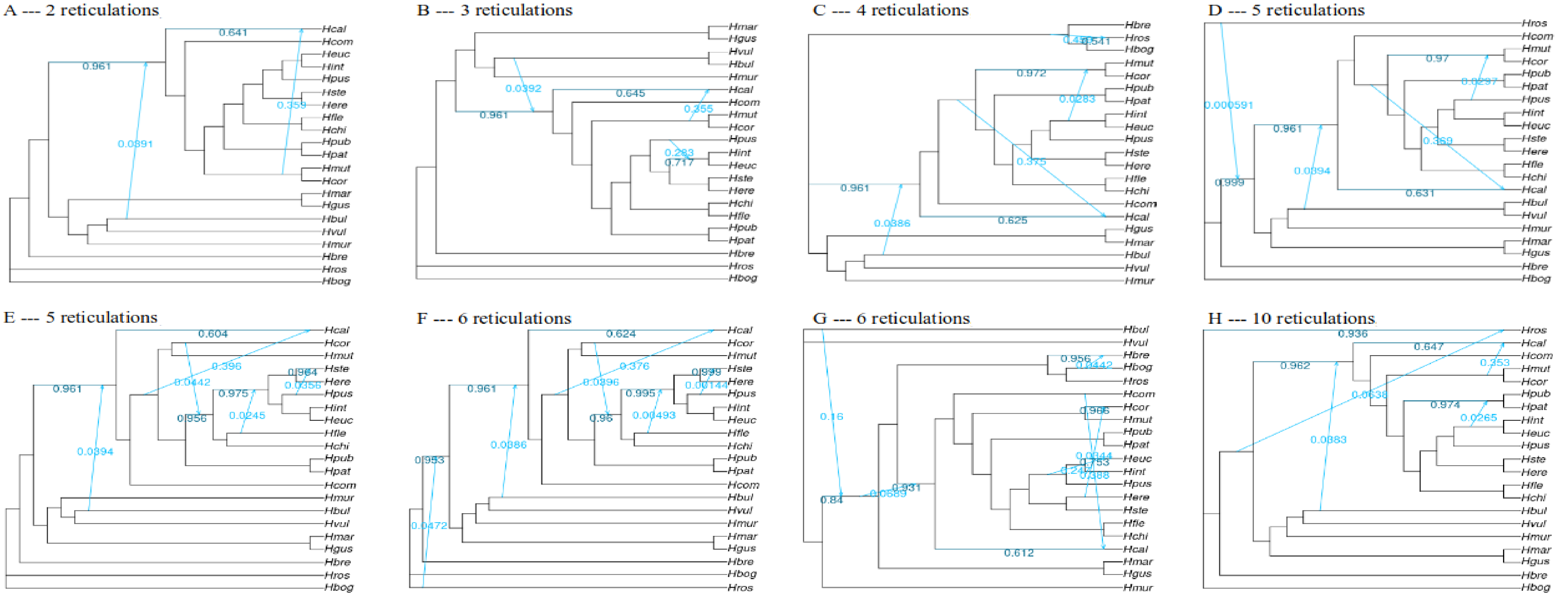


**Figure S5:** Candidate phylogenetic networks (A-H). Coloured lines indicate major (dark blue; $\gamma > 0.5$) and minor parent edges (light blue; $\gamma < 0.5$) of putative hybridization events. Their corresponding numbers represent the inheritance proportion ($\gamma$).

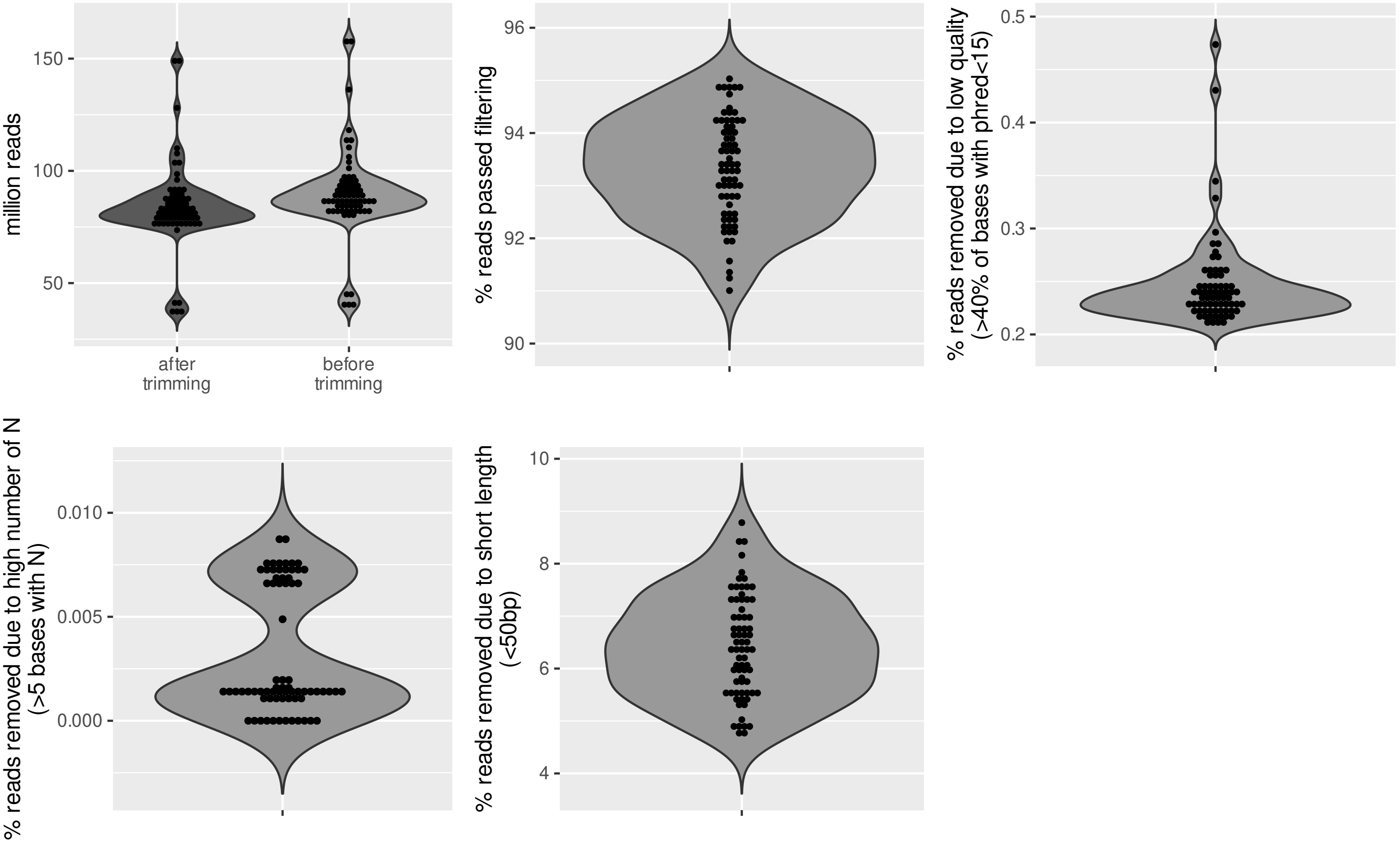


**Figure S6:** Summary statistics of the read filtering process. Each dot represents one accession.

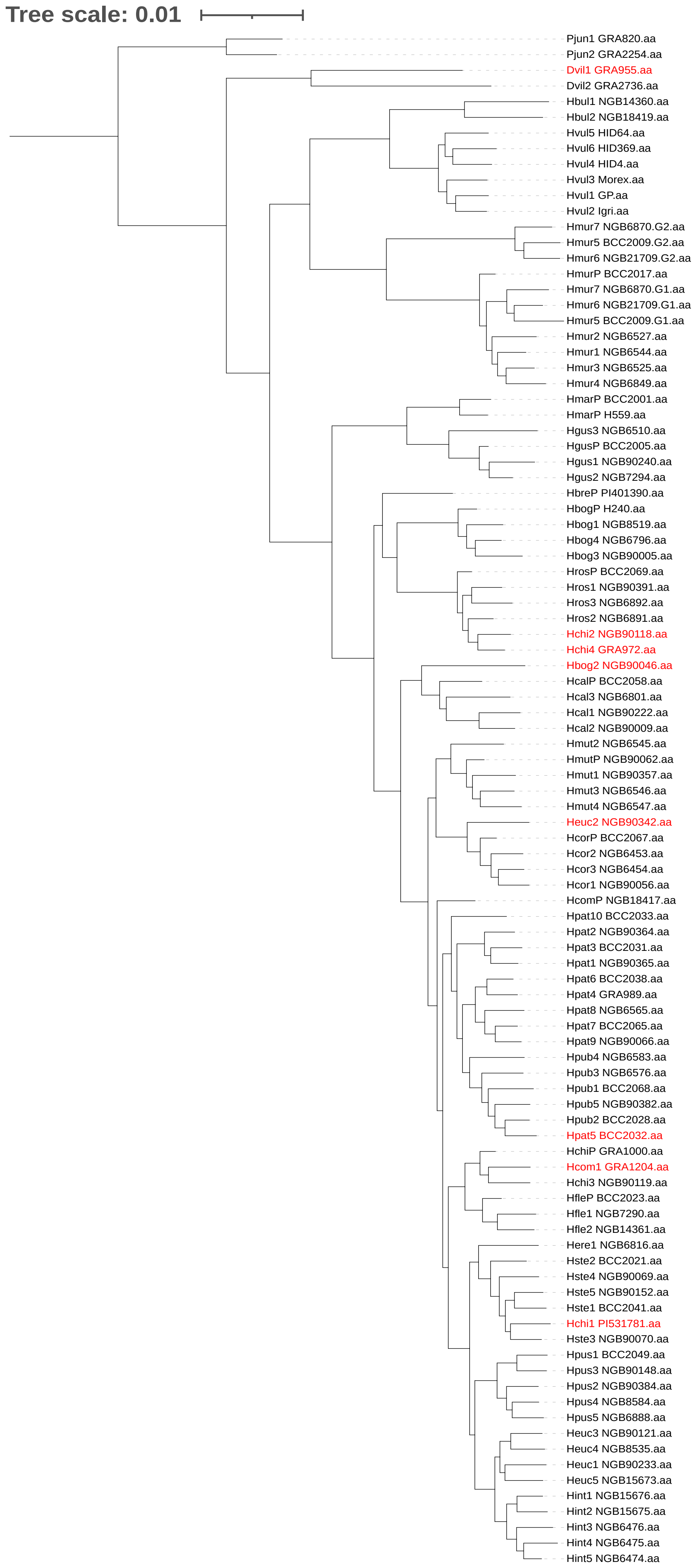


**Figure S7:** Phylogenetic tree of the unpruned set of accessions created by fasttree during the first OrthoFinder analysis. Accessions in red were removed due to their possible mislabeling.

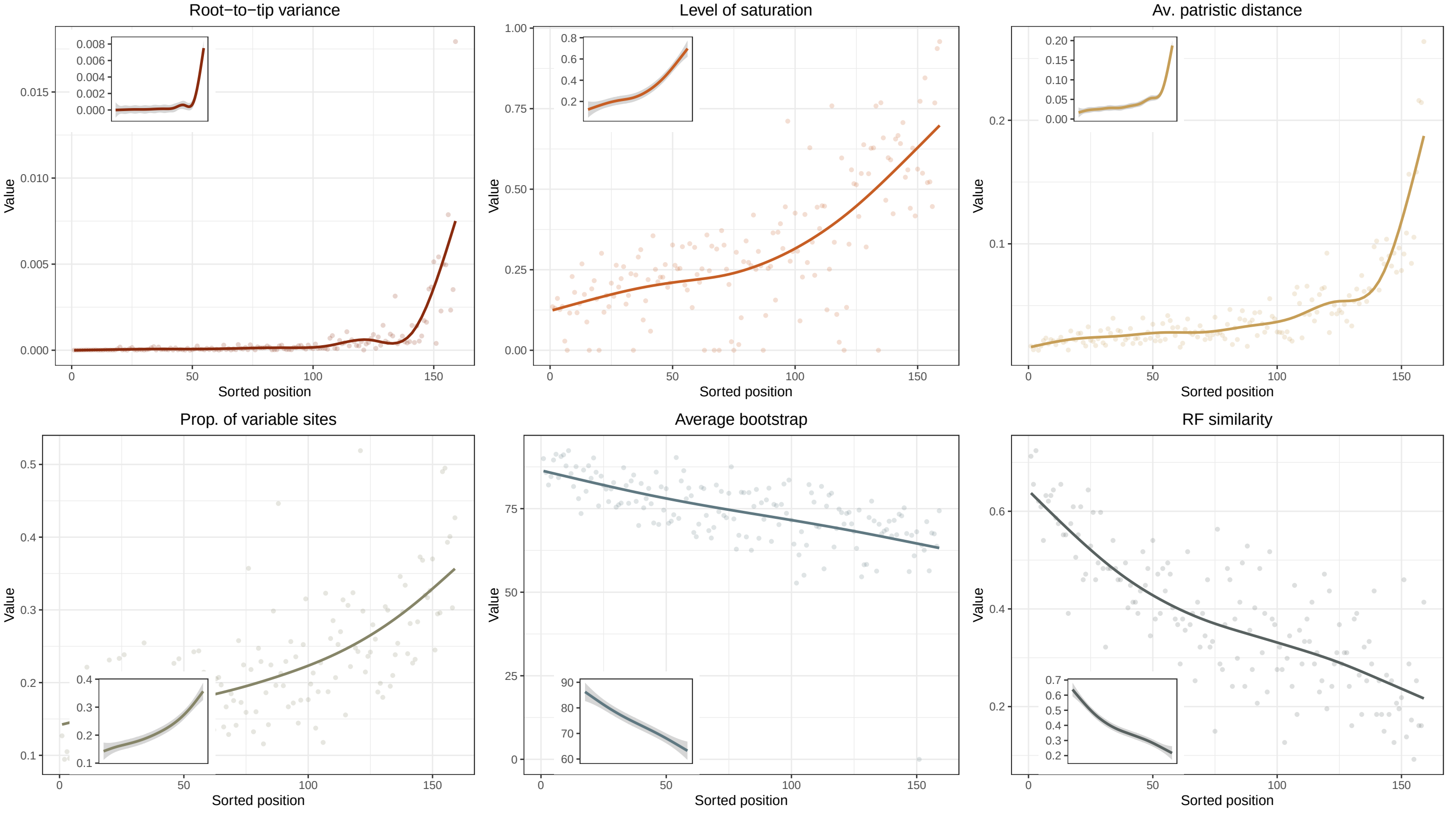


**Figure S8:** Value of the six gene-partition properties against their sorted position. Order was determined according to their phylogenetic usefulness estimated by genesortR.
